# *Cryptococcus neoformans* can form titan-like cells *in vitro* in response to multiple signals that require the activation of several transduction pathways

**DOI:** 10.1101/193540

**Authors:** Nuria Trevijano-Contador, Suelen A. Rossi, Haroldo Cesar de Oliveira, Irene Llorente, Inês Correia, Jesús Pla, Ángel Zaballos, Joaquín Ariño, Oscar Zaragoza

**Affiliations:** Mycology Reference Laboratory, National Centre for Microbiology, Instituto de Salud Carlos III, Majadahonda, Madrid, Spain.; Universidade Estadual Paulista (UNESP), Faculdade de Ciências Farmacêuticas, Câmpus Araraquara, Departamento de Análises Clínicas, Laboratório de Micologia Clínica, Araraquara, São Paulo, Brazil.; Department of Microbiology II, Complutense University of Madrid, Madrid, Spain.; Genomics Unit, Core Scientific Services, Instituto de Salud Carlos III, Majadahonda, Madrid, Spain.; Institut de Biotecnologia i Biomedicina and Departament de Bioquímica i Biologia Molecular, Universitat Autònoma de Barcelona, Spain.

**Author notes:** Present Address: Albert Einstein College of Medicine. Bronx, New York.

**Keywords:** *Cryptococcus neoformans*, titan-like cell, capsule, quorum sensing, RNAseq

## Abstract

*Cryptococcus neoformans* is an encapsulated pathogenic yeast that can change the size of the cells during infection. In particular, this process can occur by enlarging the size of the capsule without modifying the size of the cell body, or by increasing the diameter of the cell body, which is normally accompanied by an increase of the capsule too. This last process leads to the formation of cells of an abnormal enlarged size denominated titan cells. Previous works characterized titan cell formation during pulmonary infection but research on this topic has been hampered due to the difficulty to obtain them *in vitro*. In this work, we describe *in vitro* conditions (low nutrient, serum supplemented medium at neutral pH) that promote the transition from regular to titan-like cells. Moreover, addition of azide and static incubation of the cultures in a CO_2_ enriched atmosphere favored cellular enlargement. This transition occurred at low cell densities, suggesting that the process was regulated by quorum sensing molecules and was independent of the cryptococcal serotype/species. Transition to titan-like cell formation was impaired by pharmacological inhibition of PKC and TOR signaling pathways. Mutants affected in capsule synthesis did not form titan-like cells. Analysis of the gene expression profile in titan-like cells indicated that they overexpressed membrane proteins and transporters, being the gene encoding the Cig1 mannoprotein involved in iron uptake from heme groups the gene most differentially expressed compared to cells of regular size. We also investigated the gene expression profile of titan-like cells isolated from mice, and observed that during infection these cells mainly overexpressed genes related to metabolism and respiration. In summary, our work provides a new alternative method to investigate titan cell formation devoid the bioethical problems that involve animal experimentation.

**AUTHOR SUMMARY:** *Cryptococcus neoformans* is a fungal pathogen that has a significant incidence in HIV+ patients in particular, in Subsaharian Africa, Asia and South America. This yeast poses an excellent model to investigate fungal virulence because it develops many strategies to adapt to the host and evade the immune response. One of the adaptation mechanisms involves the formation of Titan Cells, which are yeast of an abnormal large size. However, research on these cells has been limited to in vivo studies (mainly in mice) because they were not reproducibly found in vitro. In this work, we describe several conditions that induce the appearance of cells that mimic titan cells, and that we denominated as titan-like cells. The main factor that induced titan-like cells was the addition of serum to nutrient limited media. This has allowed to easily performing new approaches to characterize several signaling pathways involved in their development. We found that the formation of these cells is regulated by quorum sensing molecules, and that pathways such as PKC and TOR kinases regulate the process of cellular enlargement. We have also to perform transcriptomic analysis, which led to the identification of new genes that could be involved in the process. This work will open different research lines that will contribute to the elucidation of the role of these cells during infection and on the development of cryptococcal disease.

## INTRODUCTION

*Cryptococcus neoformans* is a basidiomycetes yeast widely distributed in the environment that can behave as a pathogen in susceptible patients [1, 2]. *Cryptococcus neoformans* can survive in the lung, but in immunosuppressed patients it can also spread to the central nervous system and cause meningoencephalitis [2]. Cryptococcal infections are major causes of death in HIV patients. Although the incidence has significantly decreased in developed countries due to the introduction of antiretroviral therapy (ART), associated mortality remains high [3, 4]. Moreover, infections by this yeast still present a high incidence in developing areas, such as the sub-Saharan Africa and Southeast Asian [5, 6].

One of the most characteristic features of *Cryptococcus* is its ability to adapt to the lung environment and to evade the host immune response. Several factors contributing to cryptococcal adaptation to the lung have been described. The most important is the presence of a polysaccharide capsule [7-9], which is antiphagocytic and protects the yeasts from stress conditions [2, 10]. The size of the capsule is not constant, and it increases during the first hours of interaction with the host [11], which indicates that this process is a response that contributes to immune evasion. Furthermore, the capsular polysaccharide is also secreted to the extracellular media where it induces immunological paralysis through multiple mechanisms [9, 12-15]. *Cryptococcus neoformans* is also a facultative intracellular pathogen in phagocytic cells [16-19], which is another important factor that contributes to fungal survival in the host. *Cryptococcus* has also developed other adaptation mechanisms that contribute to the evasion of the immune response. One of them involves the formation of titan cells, which have an abnormal large size. The regular diameter of *Cryptococcus* cells grown *in vitro* ranges between 4-7 microns. In contrast, the fungal population in the lungs is very heterogeneous, and cells of even 100 microns have been described [11, 20-22]. Titan cells have been arbitrarily defined as those with a cell body diameter above 15 microns or with a total size (capsule included) over 30 microns [23]. Because of their size, titan cells cannot be phagocytosed, and they can persist in the host for long periods [24, 25]. Titan cells also contribute to virulence through other mechanisms. For example, they can divide and produce a progeny of regular size that has increased resistance to stress factors [26] as well as to inhibit the phagocytosis of cells of regular size [25].

Some signals and pathways involved in titan cell formation have been characterized, and it is known that the cAMP pathway is required for this phenomenon [20, 21, 27]. Titan cell formation has been associated *in vivo* with anti-inflammatory Th2 type immune responses [23], but the host’s factors that trigger this morphological transition remain unknown

The investigation and characterization of titan cells has been limited by the lack of media allowing the transition *in vitro*, so most of the data about these cells has been obtained using animal models. Although this approach has been shown to be useful for some purposes, it has the limitation to obtain large population of titan cells. In addition, the use of mice for these purposes presents associated significant bioethical issues. In this work, we have defined *in vitro* conditions that induce cell enlargement in *C. neoformans*, which lead to the appearance of cells similar to those found *in vivo,* which we denominate *titan-like cells*. We found that incubation of this fungus in low nutrient media supplemented with serum in a CO_2_ enriched atmosphere induced cryptococcal cell size increase. Moreover, other factors, such as oxygen limitation, or low cell density enhanced the cell growth. We have used this medium as a first step in the characterization of this transition. Our findings open future research lines that will help to define the molecular mechanisms that trigger titan cell formation and their role during infection.

## RESULTS

### *In vitro* titan-like cells formation of *C. neoformans*

In the last years, we have characterized the phenomenon of capsule growth in vitro using a medium that contains 10% Sabouraud buffered at pH 7.3 with 50 mM MOPS. In a set of experiments, we observed that addition of serum (FBS) and of the respiration inhibitor sodium azide to this medium produced not only growth of the capsule but also of the cell body (Fig 1A-C). These cells resembled the titan cells observed *in vivo*, although they did not reach the same size. Despite this difference, we argued that this phenomenon might reflect the first steps of titan cell formation, so we decided to characterize the factors that induced *in vitro* cell growth and denominated these cells as “*titan-like cells”*. First, the morphology of *C. neoformans* was analyzed in regular growth conditions (liquid Sabouraud), in capsule inducing medium (10% Sabouraud buffered with 50 mM MOPS) and in 5% Sabouraud buffered with MOPS and 5% FBS + azide (15 μM). Titan cells have been defined as those with total diameter of 30 μm or those with a cell body diameter larger than 15 μm [22]. As shown in Fig 1D, we observed a significant increase of both the cell size and the capsule in this last medium (p<0.05), almost reaching the threshold for titan cells definition after three days of culture.

**Figure 1:**
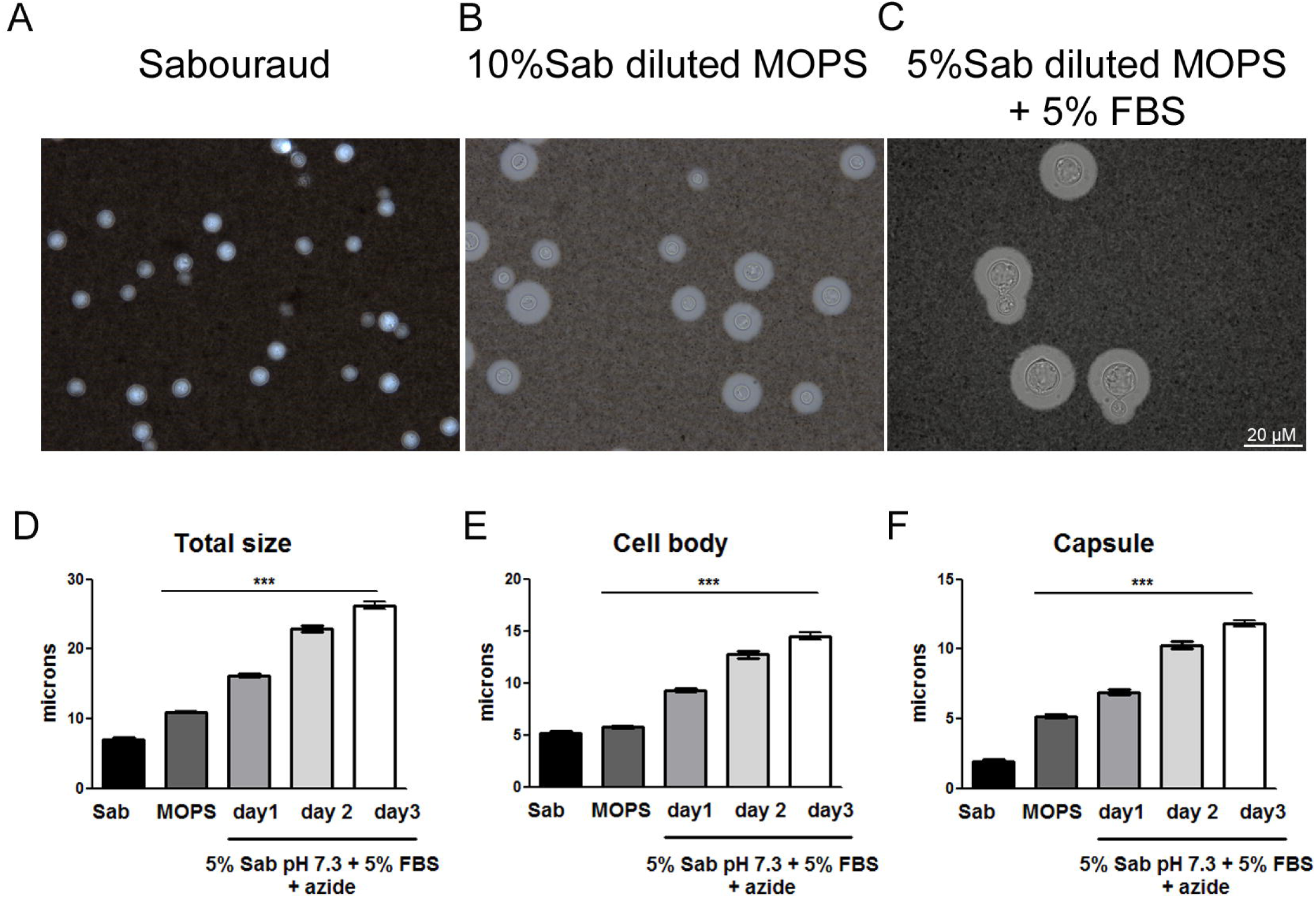
Cellular and capsular size of *C. neoformans* in different media. Cells from strain H99 were inoculated into Sabouraud (A), capsule inducing medium (10% Sabouraud with MOPS at pH 7.3) (B) and 5% Sabouraud with MOPS + 5% FBS + sodium azide (C). After incubation at 37 °C with shaking, total size and capsule size was visualized by suspending the cells in India ink. Total size (D), cell body size (E) and capsule size (F) distribution after incubation as described above. The asterisks indicate significant differences compared to Sabouraud control (p<0.05). Sab, Sabouraud medium

### Characterization of factors that induce titan-like cells *in vitro*

Serum was essential for titan-like cells formation because in its absence the increase in cell size was significantly lower (p<0.05, Fig 2A). We wanted to characterize in detail the factors and conditions that favor cryptococcal cell size increase *in vitro*. In our initial experiments, the medium in which we first observed titan-like cells contained subinhibitory concentrations of the mitochondrial inhibitor sodium azide to prevent contamination. As shown in Fig 2B, the process of cell enlargement was enhanced in the presence of sodium azide. This compound is an inhibitor of complex IV of the respiratory chain, so we argued that titan-like cell formation could be induced by factors that alter respiration. For this reason, we investigated if titan-like cell formation was influenced by oxygen limitation. We compared the formation of titan-like cells under shaking or static conditions. As shown in Fig 2C, static incubation of the cultures resulted in a greater proportion of titan-like cells.

**Figure 2:**
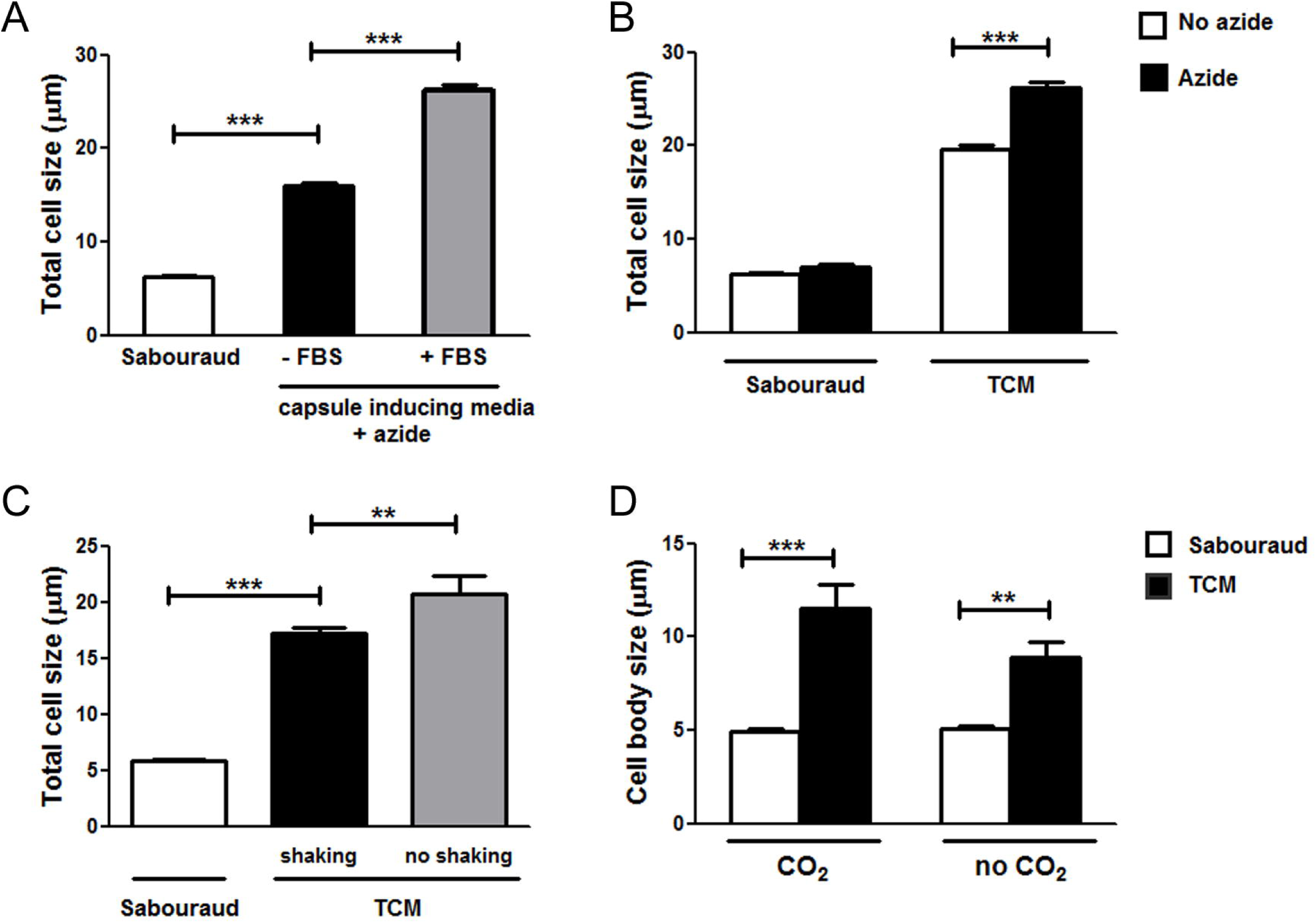
Effect of different factors on the cellular growth of *C. neoformans*. Cells from H99 strain were incubated on Sabouraud overnight and transferred to different media to evaluate the effect of several factors on the titan-like cell formation. (A) Cells were incubated in Sabouraud or capsule inducing medium (10% Sab, pH 7.3) supplemented with 15 μM sodium azide in the presence or absence of 5% serum (FBS) and cultures were incubated at 37 °C overnight. Pictures after suspension of the cells in India Ink were taken, and the total cell size was measured and plotted (B) Effect of sodium azide (black bars) on cell size in Sabouraud or TCM medium. As a control, the same media were inoculated without sodium azide (white bars). Cells were incubated at 37 °C overnight. Pictures after suspension of the cells in India Ink were taken, and the total cell size was measured and plotted (C) Effect of shaking effect on titan-like cells formation. The yeasts were incubated at 37°C for 24 h in flasks with Sabouraud or TCM medium in both conditions (black bars, shaking) and (gray bars, no shaking). Pictures after suspension of the cells in India Ink were taken, and the total cell size was measured and plotted (D) Effect of CO_2_ on cell growth. The cells were incubated in Sabouraud (white bars) and TCM (black bars) with and without 5% of CO_2_. In both cases, the cells were grown at 37°C without shaking for 24 h in a 96-microdilution plate. Then, the plate was directly observed in the microscope, and the cell body size of was measured and plotted.

Cryptococcal cells sense and respond to environmental levels of CO_2_ and it is known that this molecule induces capsule growth. For this reason, we investigated if incubation of the cultures in a CO_2_-enriched environment altered the formation of titan-like cells. We found that the induction of these cells was enhanced when the plates were placed in a 5% CO_2_ atmosphere in comparison to growth without CO_2_ (Fig 2D).

Although serum was required to obtain titan-like cells *in vitro*, it was not sufficient for titan-like cell formation because incubation of the cells in 100% serum did not result in cellular growth (result not shown). Furthermore, serum did not induce cellular growth in rich media (Fig 3A and B), in contrast to the situation in low nutrients medium (figure 3C), indicating that nutrient limitation was important for titan-like cell formation.

**Figure 3:**
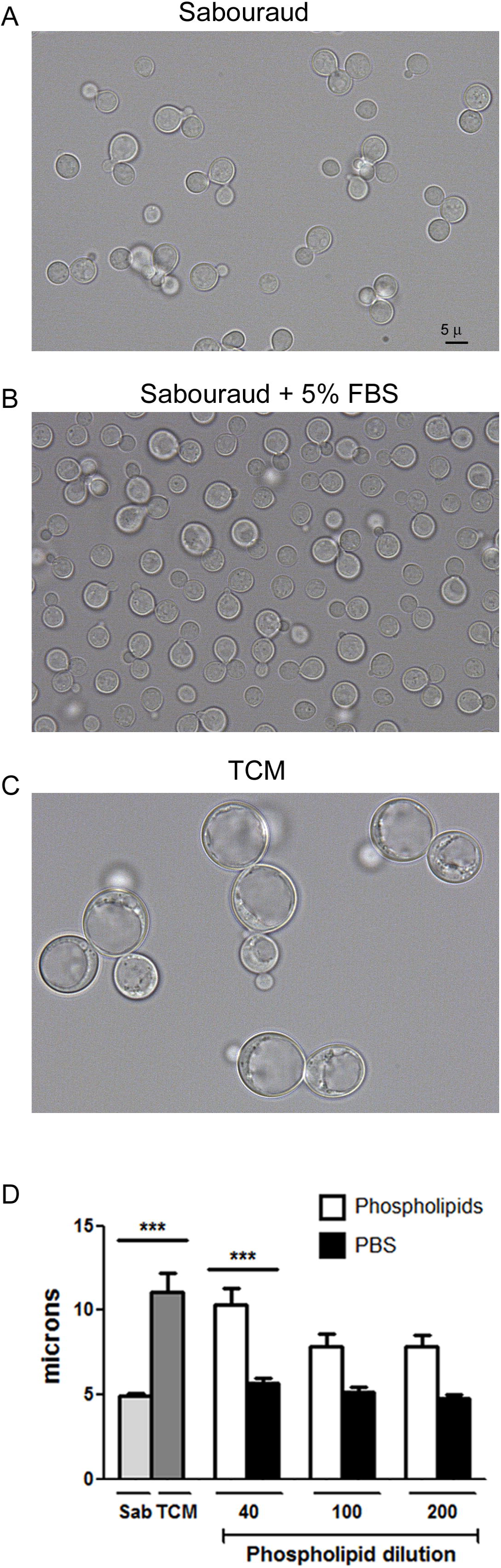
Effect of serum on cryptococcal cell growth. H99 cell suspensions were inoculated in Sabouraud (A), Sabouraud supplemented with 5% serum (B) or TCM (C). The cells were incubated overnight at 37 °C in 5% CO_2_. Then, the cells were suspended in India Ink and pictures were taken. The scale in A applies to all the pictures. (D) Effect of purified serum lipids on cell growth. The cells (initial density 10^4^ cells/mL) were grown in TCM medium in which the serum was replaced by different dilutions (40, 100 and 200) of the lipid extract. As controls, the same media was supplemented with PBS. Cultures in TCM are also included for reference. Cell body size was determined after 24 hours at 37° C with CO_2_ and shaking. The asterisks indicate significant differences compared to Sabouraud control.

To visualize the phenomenon of titan-like cell formation, we carried out *in vivo* imaging by placing the cells in a 96-wells plate in TCM in 5% CO_2_ at 37 °C in a microscope overnight and obtained videos of the cellular enlargement. As shown in supplemental video 1, cells actively grew and replicated in Sabouraud medium. However, in TCM, after 5-8 h of incubation the cells started to enlarge during 8-10 h (supplemental video 2). After this time, the cells stopped growth and continued budding. We also observed that titan-like cell formation was associated with some intracellular features. We visualized in a significant proportion of the cells that there was an intracellular compartment that started to divide by fission, but then fused again to render a large vesicle (Supplemental video 3).

### Effects of phospholipids from fetal calf serum in titan-like cells formation

Phospholipids, in particular phosphatidylcholine, can trigger the appearance of titan cells *in vitro* [28]. For this reason, we performed a lipid extraction of fetal calf serum, which is present in our medium (TCM) and we incubated the cells with different amounts of these lipids (1/40, 1/100 and 1/200 dilution of the original lipid solution). As shown in Fig 3D, the lipids present in the serum induced titan-like cell formation in comparison with the same amount of PBS.

### Cell density influences titan-like cells formation

We found that titan-like cell formation depended on the cell density of the cultures. We inoculated 96-wells plates with different concentrations of cells from H99 strain (10^6^, 10^5^, 10^4^ and 10^3^ cells/mL) in TCM and Sabouraud as a control of growth. After overnight incubation at 37 °C with CO_2_, titan-like cells were observed in the wells inoculated with 10^3^, 10^4^ and 10^5^ cells/mL but were almost absent in the wells that were inoculated with the higher cell density (10^6^ cells/mL, Fig 4A). We quantified the morphological difference by measuring the cell body of 50-60 cells per condition. We found that titan-like cells were observed more frequently when the cultures were inoculated with cellular concentrations around 10^4^ cells/mL.

**Figure 4:**
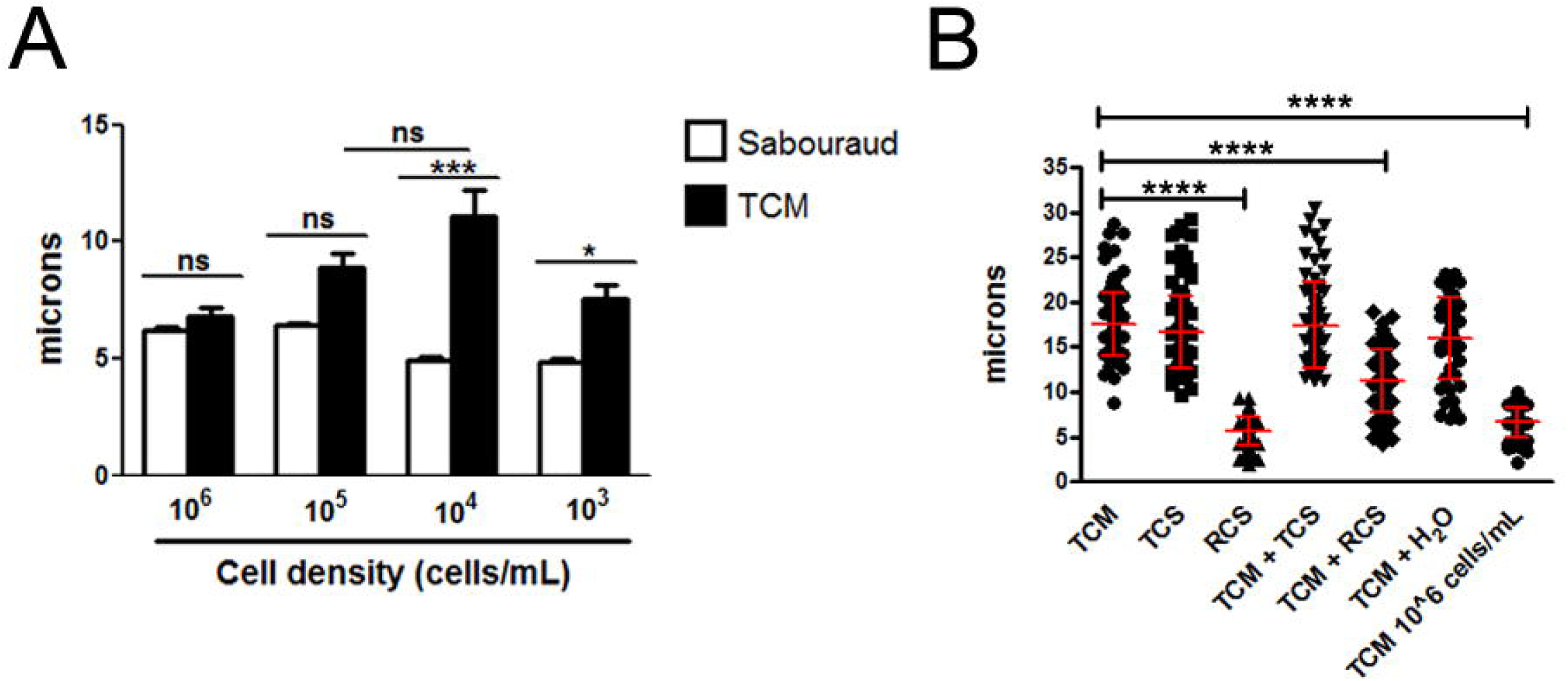
Influence of cell density on titan-like cell formation in. ***C. neoformans***. (A). Cells from H99 strain inoculated in Sabouraud or TCM medium at different concentrations (10^3^, 10^4^, 10^5^ and 10^6^ cells/mL) in 96-well plates. Cell body size was measured after incubation overnight at 37 °C with CO_2._ The asterisks indicate significant differences. B) Effect of conditioned media on cell body size. Cells from H99 strain were incubated in different conditioned TCM medium denominated (see material and methods) as TCS (supernatant of titan-like cells TCM cultures), RCS (supernatant of regular size TCM cultures), TCM + TCS (a 1:1 mixture of fresh TCM with TCS), TCM + RCS (a 1:1 mixture of fresh TCM with RCS). As controls we used a diluted TCM in H_2_O (1:1) and fresh TCM inoculated with 10^4^ cells/mL (TCM) and with 10^6^ cells/mL.

### Effect of QS molecules on titan-like cell formation

The fact that the formation of titan-like cells depends on cell density suggests that this process can be regulated by quorum sensing (QS). QS is a cell-cell communication mechanism mediated by molecules that are released directly into the medium by microorganisms. These molecules are released as a function of growth and replication rate [29, 30]. In this way, we evaluated the influence of titan-like and regular *C. neoformans* supernatants cultures on titan-like cell formation. We inoculated TCM with the H99 strain at 10^6^ cells/mL and 10^4^ cells/mL and incubated the cultures for 18 h at 37 °C in 5% CO_2_ to obtain cells of regular size and titan-like cells, respectively. We then collected the supernatants (named RCS and TCS, respectively). These conditioned media were added to wells that contained fresh TCM inoculated at 10^4^ cells/mL. We found that the conditioned medium RCS significantly (p<0.001) inhibited the formation of titan-like cells (Fig 4B) even when added to fresh TCM (TCM + RCS) (p<0.001). In contrast, the supernatant from titan-like cells cultures (TCS) did not block the formation of the titan-like cells, demonstrating a negative effect on titan-like cells formation of the supernatant obtained from cells of regular sizes (Fig 4). The effect of the TCS conditioned medium was not explained by a the dilution of the nutrients of the fresh TCM, since titan-like cells were still formed in TCM diluted with distilled water (Fig 4B).

In *C. neoformans*, the main QS molecule described is a short peptide of 11-mer called Qsp1 which is required for fungal virulence, replication, cell wall synthesis and protease activities [31, 32]. To investigate the influence of the Qsp1 in the formation of titan-like cells, different concentrations of the peptide were added to the TCM medium and the formation of titan-like cells was evaluated. We observed that Qsp1 significantly inhibited formation of titan-like cells in a dose-dependent manner (p<0.001, Fig 5A). As control, we used both inactive and scrambled versions of Qsp1, and observed that none of them had any effect on titan-like cell development (Fig 5A).

**Figure 5:**
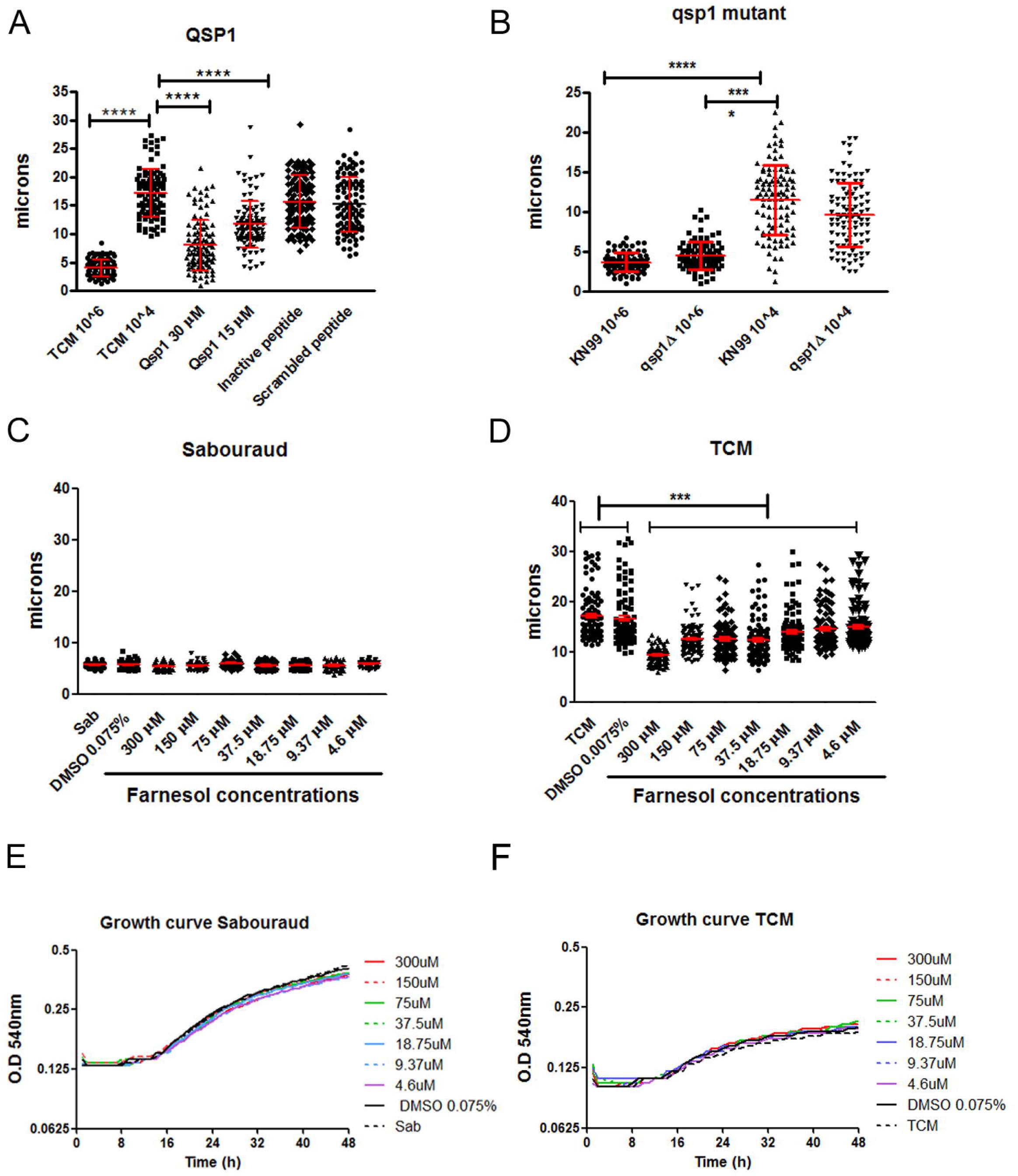
Influence of quorum sensing molecules on titan-like cell formation. A) Cells from H99 strain were grown in TCM supplemented with 30 uM and 15 uM of the quorum sensing peptide Qsp1 (NFGAPGGAYPW). As controls, TCM supplemented with 30 uM of an inactive Qsp1 (NFGAPGAAYPW), with 30 uM of a scrambled Qsp1 peptide (AWAGYFPGPNG). TCM without any supplementation was used as control. Cell body size was measured after overnight incubation at 37 °C with 5% CO_2_. B) Titan-like cell formation of *qsp1* mutant. Cells from the KN99 (wild type) and *qsp1* mutant were inoculated in TCM at 10^4^ and 10^6^ cells/mL, and cell body size was determined after 18 h of incubation at 37 °C in the presence of 5% CO_2_. C-F) Effect of Farnesol on the titan-like cell formation. Cells of strain H99 were grown on Sabouraud (C) and TCM medium (D) with different concentrations of farnesol in 96-well plates. As controls, we used media with 0.075% of DMSO (Farnesol solvent), Sabouraud and TCM medium without DMSO or farnesol. The plate was incubated at 37°C with 5% CO_2_ overnight without shaking and cell body size was measured. The asterisks indicate significant differences (p<0.05). Growth curves of strain H99 with different concentrations of farnesol in Sabouraud (E) and TCM (F) without shaking. Cells were incubated at a density of 2x10^5^ cells/mL during 48 hours at 37 °C, measuring the optical density at 540 nm.

It could be argued that the production of Qsp1 in TCM cultures inoculated at high cell densities was responsible for the inhibition of titan-like cells formation. To test this idea, we used a *qsp1* mutant that does not produce Qsp1 [32]. Our results showed that the mutant produced titan-like cells in a similar way as the wild type strain KN99 (Fig 5B) [32], even at high cell densities (10^6^ cells/mL). This result indicates that absence of titan-like cells in TCM cultures inoculated at high densities was not only due to Qsp1, and that most probably, other QS molecules secreted by *C. neoformans* might influence cellular enlargement.

Farnesol is a sesquiterpene alcohol and the first QS molecule described in eukaryotes. Although *C. neoformans* does not produce this compound, we thought that it would be interesting to test if this yeast responded to QS molecules produced by other microorganisms. We evaluated the effect of Farnesol (300 μM to 0.5 μM) on the formation of titan-like cells in Sabouraud and TCM in microdilution plates. We observed that Farnesol had no effect on cell size when grown in Sabouraud medium (Fig 5C). However, Farnesol inhibited the formation of titan-like cells in TCM medium in a dose dependent manner (5D). This inhibition did not correlate with any growth defect due to the presence of farnesol (Fig 5E,F).

### Nuclear staining

Titan cells formed in the lungs are polyploid and single-nucleated. So we investigated the morphology of the nucleus and the DNA content after staining with DAPI. As shown in Fig 6A-B, titan-like cells contained one nucleus. This result was confirmed using a strain that expresses a fluorescent nucleolar protein (NOP1-mCherry, [33], supplemental Fig 1). We also quantified the fluorescence intensity of the DAPI staining by flow cytometry. As shown in Fig 6C-E, titan-like cells emitted more fluorescence than cells of regular size. The fluorescence intensity was more heterogeneous in titan-like cells, ranging from 2 to 5 fold increase compared to cells of normal size (Fig 6E).

**Figure 6:**
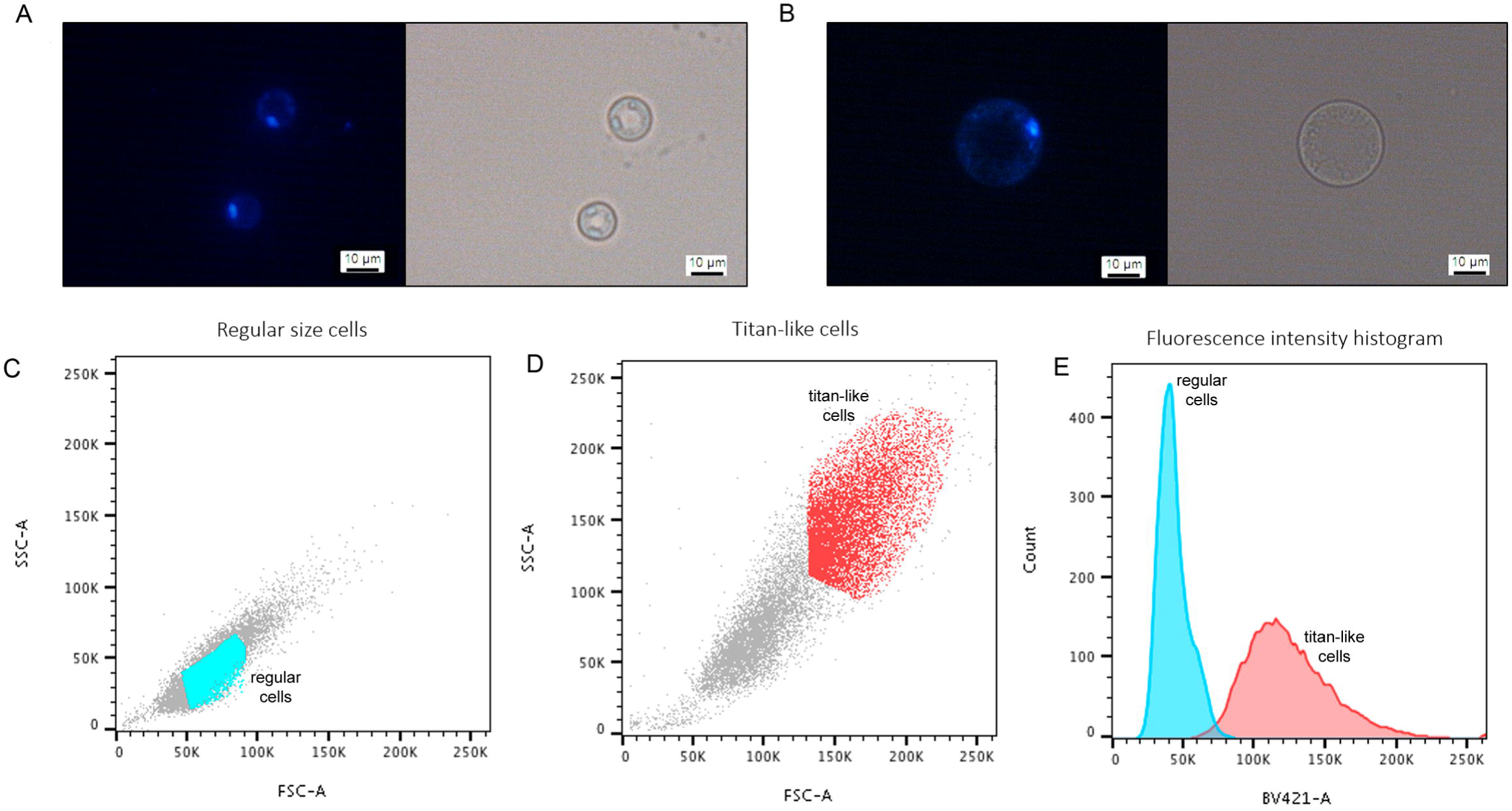
Nuclear staining of titan-like cells. Cells of regular size and titan-like cells were obtained by inoculating H99 strain in TCM at 10^6^ and 10^4^ cells/mL respectively. After incubating overnight at 37 °C with 5% CO_2_, the cells were fixed and stained with DAPI as described in material and methods. A and B show the microscopic appearance of the nucleus of cells of regular size (A), and titan-like cells (B). C-E, Analysis of nuclear staining by flow cytometry. C) FFS/SSC scatter plot of cells of regular size, D) same graph of titan-like cells. In C and E, we defined a gate to clearly separate cells of regular size (cells in blue) and titan-like cells (cells in light red). The fluorescence intensity (histograms) of the cells from these two gates is represented in panel E). Blue histogram, fluorescence from the cells shown in the gate in panel C, and light red histogram, fluorescence from the cells shown in the gate in panel D.

### Inhibition of PKC and TOR signaling pathways affected titan-like cell formation

PKC is a family of protein kinases, activated by Ca^2^+- diacylglycerol (DAG) and phospholipids, and involved in different virulence events in *C. neoformans* such as melanin production [34], temperature tolerance, cell integrity [35] and fluconazole tolerance [36]. We argued that serum phospholipids could activate the PKC signaling pathway, so we tested the effect of three PKC inhibitors, calphostin C, staurosporine and bisindolylmaleimide I on titan-like cell formation. We observed that all PKC inhibitors (in particular, staurosporine and Calphostine C), impaired the formation of titan-like cells in a dose-dependent manner (Fig 7A-C). We also included the tyrosine kinase inhibitor genistein and found that this compound had no visible effect on cellular enlargement (Fig 7D).

**Figure 7:**
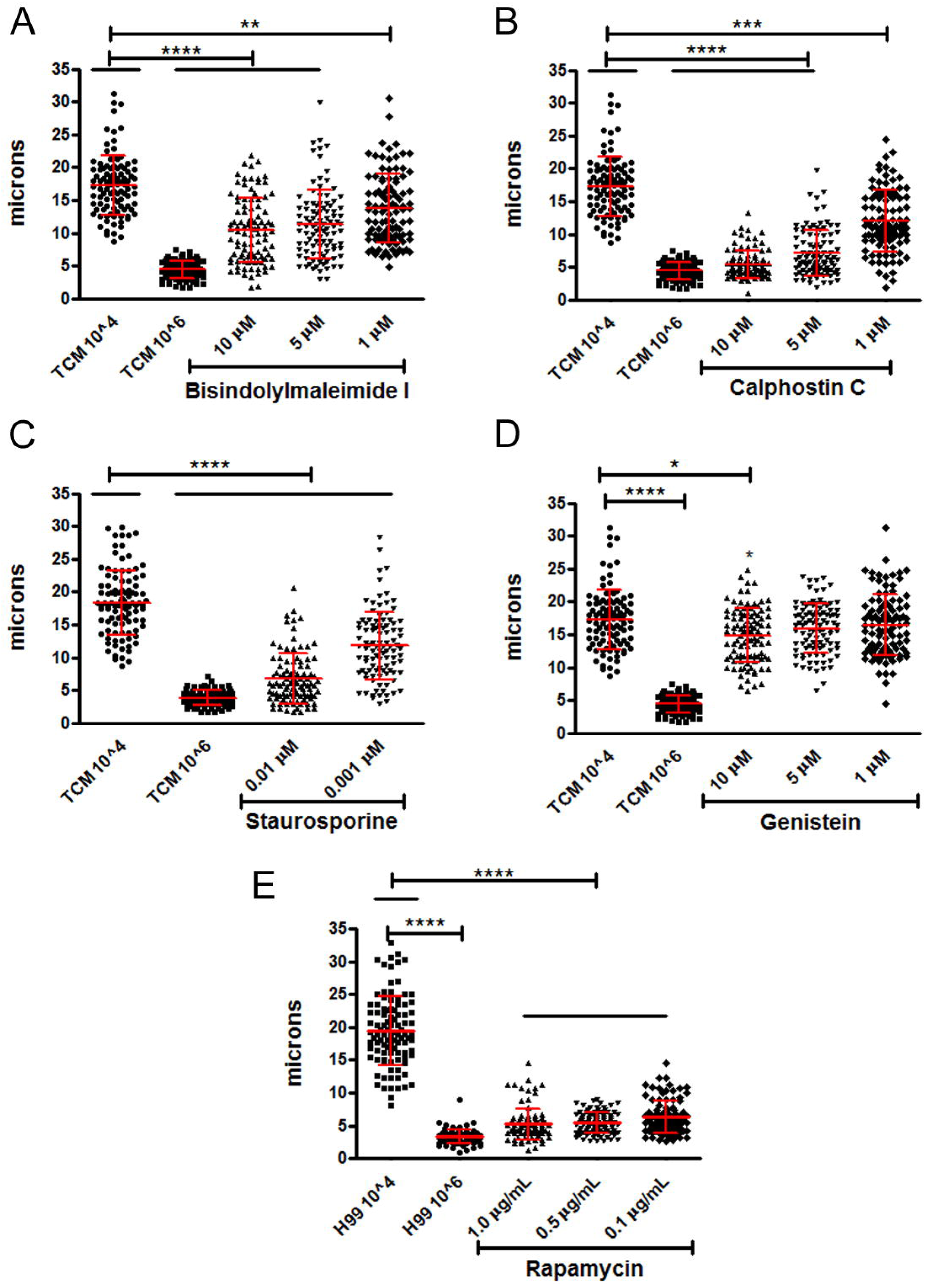
Influence of PKC and TOR pathways on titan-like cells formation. The influence of the PKC pathway was evaluated by its inhibition with three different agents: Bisindolylmaleimide I (A), Calphostin C (B), Staurosporine (C). Genestein, a tyrosine kinase inhibitor, was used as control (D). The experiments in A, B and D were performed the same days, so they share the same control. However, for clarity, the graphs corresponding to each inhibitor have been separated. The experiments were repeated on three days, and the data from the three experiments is plotted. (E) The inhibition of the TOR pathway was made with the addition of Rapamycin. The experiments (A-E) were repeated on three days, and the data from the three experiments is plotted. Cells of the H99 strain were incubated with these different agents overnight at 37 °C with 5% CO_2_ without shaking and cell body size was measured. The asterisks indicate significant differences.

We also tested the role of the TOR signaling pathway. TOR proteins are kinases that regulate cell size and replication in response to the availability of nutrients in the medium. To investigate the role of this pathway on titan-like cell formation, we inhibited it with rapamycin. As shown in Fig 7E, this compound blocked the appearance of titan-like cells in TCM. Therefore, both PKC and TOR pathways might be relevant for titan-like cell formation.

### Titan-like cell formation did not correlate with the serotype of the strain

*Cryptococcus neoformans* is divided in different serotypes and varieties: variety *grubii* (serotype A), variety *neoformans* (serotype D), and A/D hybrids. It has been proposed that these groups should named into divided species [37], although recent reports argue against this differentiation and suggest the term of species complexes [38]. We investigated if there was a correlation between the main species complex of the strain and the formation of titan-like cells. As shown in Fig 8A-C, for each serotype/species complex, there were strains with high and low capacity to induce titan-like cells, which indicates that there was not a direct association between the serotype and cell growth.

**Figure 8:**
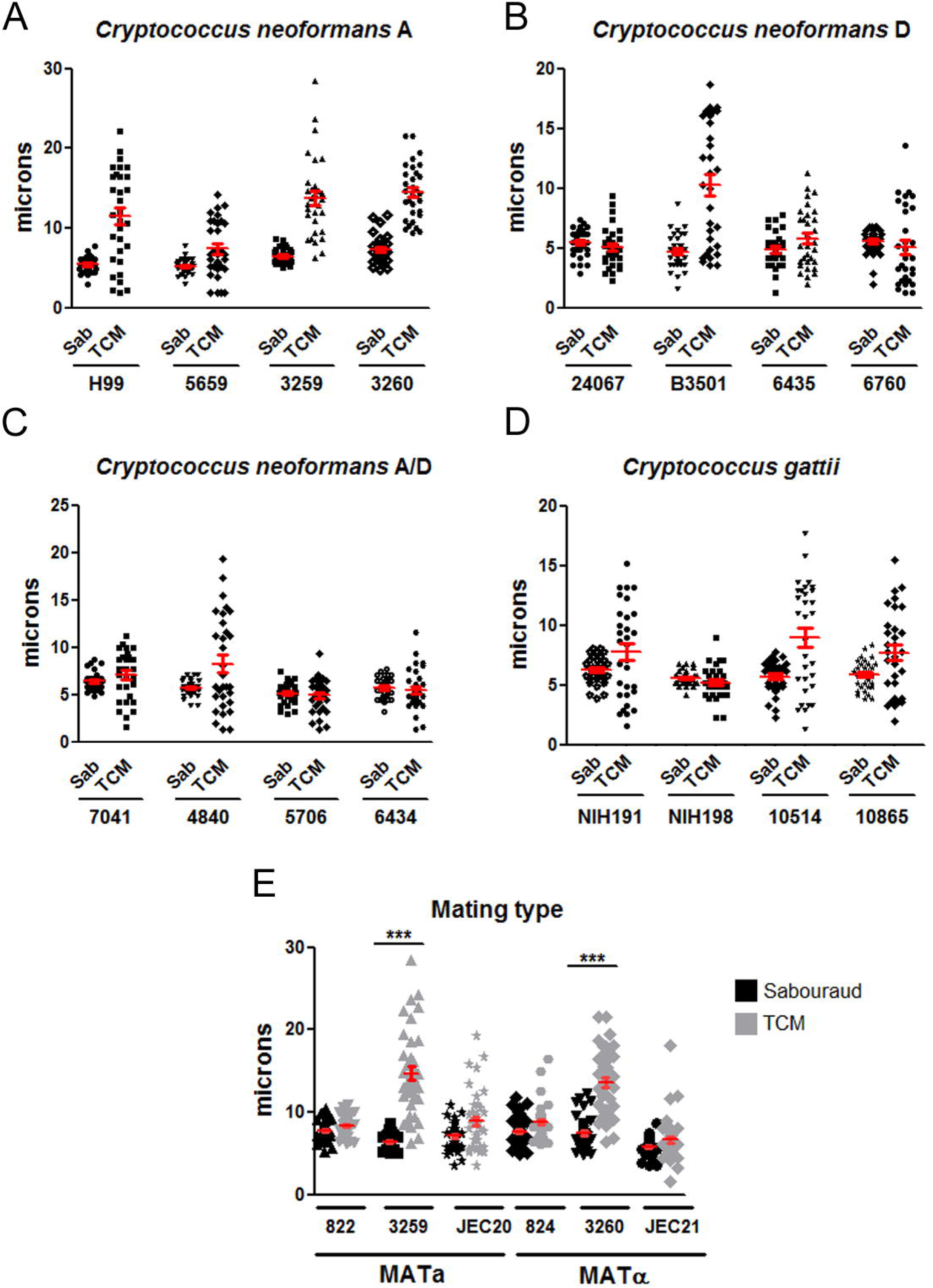
Titan-like cell induction in different strains. Four strains of each species (serotypes) were inoculated into Sabouraud or TCM medium at a density of 10^4^ cells/mL and grown overnight at 37 °C with CO_2._ *(A) C. neoformans* var. *grubii* (serotype A); (B) *C. neoformans* var *neoformans* (serotype D); (C) *C. neoformans* A/D; (D) *C. gattii*. (E) Cell body growth in MATa and MATα strains: NE822/NE824 (serotype D), 3259/3260 (serotype A) and JEC20/JEC21 (serotype D). The strains were inoculated in Sabouraud (black symbols) and medium TCM (gray symbols) and under the conditions described above. Cell body size was measured in all cases. The red lines represent the mean and standard error.

We also investigated the behavior of strains from the related species *C. gattii*, which can infect immunocompetent patients. As shown in Fig 8D, there were strains that had high and low ability to induce titan-like cells. We next examined if the hyper (CBS10514, R265) and hypovirulent (CBS10865, R272) strains isolated at the Vancouver outbreak [39, 40] formed titan-like cells in a different way. Interestingly, the hypervirulent strain tended to produce more titan-like cells compared to the strain with reduced virulence (Fig 8D). We also tested other *C. gattii* strains (NIH 191 and NIH198), which also presented difference in their capacity in produce titan-like cells. In summary, there were many inter-strain differences in their ability to form titan-like cells which were not associated to the serotype/genotype of the isolates.

### Correlation between mating type and titan-like cell formation

Coinfection with a and α strains results in a higher proportion of titan-like cells in the lungs [20], so we investigated if strains from different mating type had different ability to form titan-like cells. We studied three pair of strains with different mating type JEC20/JEC21, NE822/NE824 and 3259/3260 (KN99). We found that pair of strains 3259/3260 increased the size of the cell body in the TCM medium compared to the Sabouraud significantly (p<0.05, Fig 8E). In NE822/NE824 and JEC20/JEC21 pairs, there was a small increase in cell size in TCM and we observed a small amount of titan-like cells in this medium although this difference was not statistically significant. These results indicate that titan-like cell formation can occur *in vitro* independently of the mating type of the strains.

### Titan-like cells formation in different mutants

To identify genes that play an important role in the formation of titan-like cells, we studied the phenotype of mutants with problems to induce capsule growth, such as *gat201* or *ada2* [41, 42] or acapsular strains (*cap59* and *cap60*). As shown in Fig 9A, none of these mutants induced cellular enlargement in TCM.

**Figure 9:**
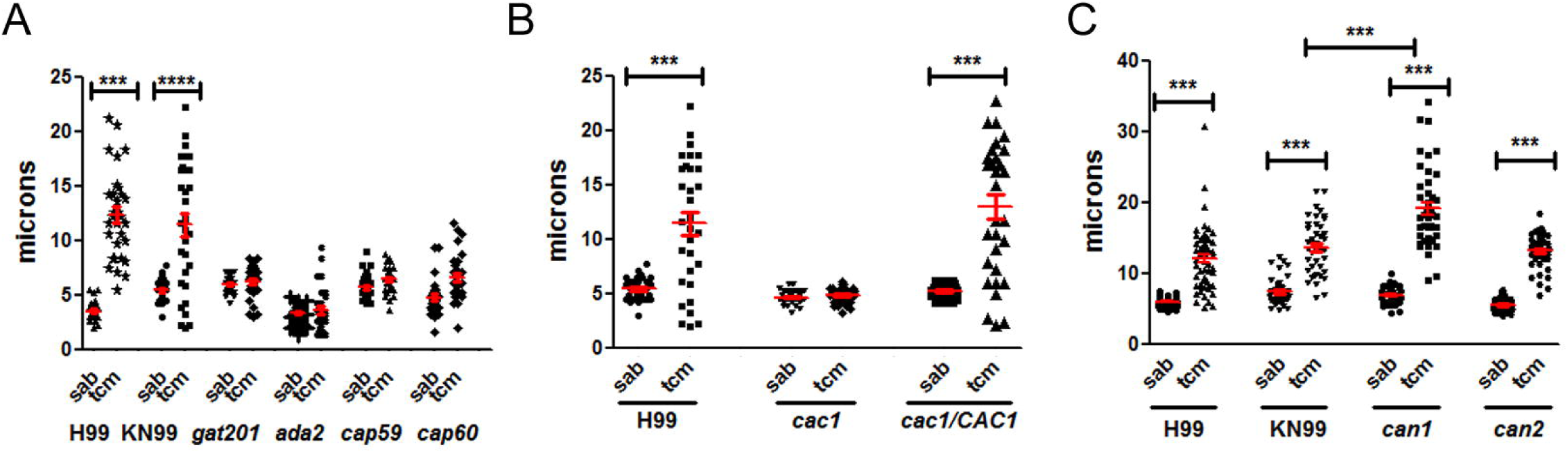
Titan-like cell formation in different mutants. (A) Cells from the capsule deficient mutants *gat201*, *cap59*, *cap60* and *ada2* (H99 background) were inoculated in Sabouraud and TCM medium at a density of 10^4^ cells/mL in 96-well plates. Cell body size was determined after overnight incubation at 37°C with CO_2._ (B) Role of adenylate cyclase on cell growth of *C. neoformans*. Cells from the wild-type strain H99, the adenylate cyclase mutant (*cac1*) and the reconstituted strain (*cac1/CAC1*) were incubated as described above. (C) Role of carbonic anhydrases on cell growth. Cells from the wild type strain H99 and KN99 and the *can1* and *can2* mutants were used for this purpose. Cell body size was determined after overnight incubation at 37 °C with CO_2_ in all cases. The asterisks indicate significant differences (p<0.05). The red lines represent the mean and standard error.

Titan cell formation is regulated by the cAMP pathway [21, 27]. Moreover, the CO_2_ activates adenylate cyclase [43, 44]. For this reason, we investigated the formation of these cells in *cac1* mutant (which lack the enzyme adenylate cyclase) and in the reconstituted strain *cac1/CAC1*. As shown in Fig 9B, the *cac1* mutant was defective to produce cellular enlargement, whereas the reconstituted strain produced titan-like cells as the wild type.

CO_2_ is transformed into HCO_3_^-^ by the action of carbonic anhydrases (Can). In *C. neoformans*, there are two genes encoding these enzymes (*CAN1* and *CAN2*) [44, 45], being Can2 the most abundant and physiologically active. Since *can2* mutant can only grow in a CO_2_ enriched environment, these strains were maintained always in 5% CO_2_. Deletion of *CAN2* did not have any effect of titan-like cell formation. Strikingly, in the absence of *CAN1*, cell size was larger than that observed for the WT strain (Fig 9C).

### Interaction of titan-like cells with macrophages

Titan c*ells are not phagocytosed in vivo* [21, 25]. For this reason, we examined the interaction between macrophage-like cell lines and titan-like cells obtained *in vitro*. We compared the phagocytosis of titan-like cells (incubated in TCM inoculated at 3x10^4^ cells/mL), and of cells of regular size obtained in TCM inoculated at 10^6^ cells/mL or in Sabouraud. Phagocytosis was quantified both by microscopic observation and by flow cytometry using a cryptococcal strain that expressed GFP (see Material and Methods). In all cases, we found that titan-like cells obtained *in vitro* were not phagocytosed (Fig 10A and supplemental video 4). Interestingly, cells of regular size were not equally internalized, since cells cultivated in Sabouraud medium inoculated at high cellular density were more efficiently phagocytosed than the same cells inoculum grown in TCM (supplemental videos 5 and 6, Fig 10A). This difference was not due to difference in binding to the antibody used as opsonin in this experiment (18B7, result not shown), but correlated with the increase in cell size (in particular, due to capsule enlargement) found in cells inoculated in TCM at high density compared to the cells cultivated in Sabouraud (data not shown).

**Figure 10:**
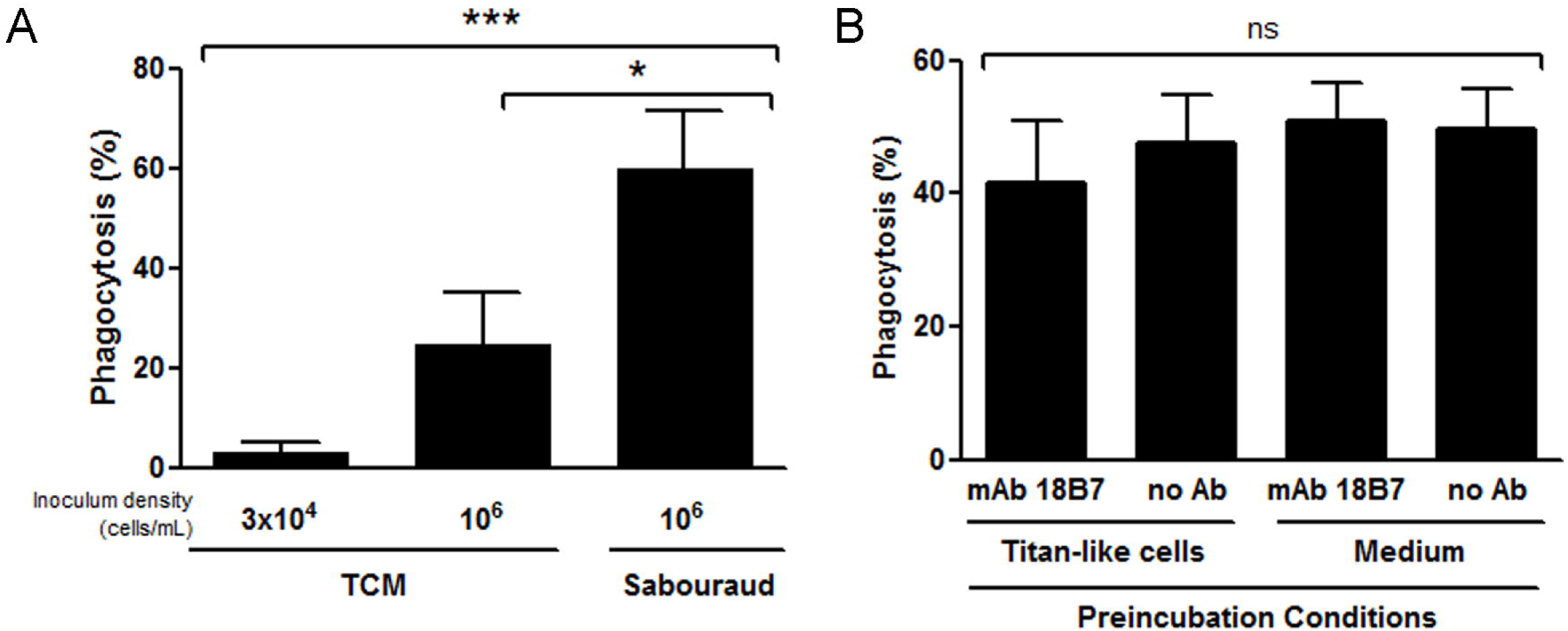
Interaction of titan-like cells with murine-like macrophages. A) Cells from H99-GFP strain were incubated in TCM inoculated at 3x10^4^ cells/mL (titan-like cells), 10^6^ cells/mL (regular cells in TCM), and Sabouraud. Then, phagocytosis experiments with RAW264.7 cells were performed and quantified by flow cytometry as described in material and methods. Statistical differences are highlighted. B) Effect of preincubation of macrophages with titan-like cells on the phagocytosis of regular cells. Titan-like cells were obtained by incubation of *C. neoformans* in TCM as described in material and methods, and were exposed to RAW264.7 macrophages at 1:1 ratio. The incubation was performed in media containing the opsonizing mAb 18B7 or in its absence. As control, macrophages were preincubated with growth medium with or without the mAb. The plates were incubated for 1h at 37 °C in the presence of 5% CO_2_, and then the cells were washed to remove titan-like cells. Next, *C. neoformans* cells (H99-GFP) of regular size grown in Sabouraud were added to the macrophages at 1:1 ratio in the presence of mAb 18B7, and phagocytosis was performed for 1 h at 37 °C in CO_2_. Phagocytosis percentage was quantified by flow cytometry as described in M&M. ns: no statistical difference. The experiment was performed in triplicates in three different days.

It has been shown that titan cells can also prevent phagocytosis *in vivo* of cryptococcal cells of regular size [25], so we tested if titan-like cells induced a similar phenomenon *in vitro*. For this purpose, RAW264.7 macrophages were preincubated with titan-like cells for 1 h with mAb 18B7. As control, parallel samples were incubated only with titan-like cells without mAb or with medium without yeast cells (with or without mAb). After this time, the plate was washed to remove titan-like cells, and H99-GFP cells of normal size cultivated in liquid Sabouraud were added to the macrophages. As shown in Fig 10B, preincubation of the macrophages with titan-like cells did not affect the phagocytosis de regular size yeasts (Fig 10B).

### Analysis of titan-like cells gene expression profiles *in vitro and in vivo*

To gain insights about the molecular mechanisms involved in titan-like cell formation, we compared their gene expression profile with that of cells of regular size. Since the number of titan-like cells obtained in TCM inoculated at low densities and grown in static conditions in the CO_2_ incubator was low, it was difficult to isolate enough RNA to perform transcriptomic profiling in these conditions. For this reason, we decided to obtain titan-like cells in TCM, but with shaking and after passaging the cells for three days to fresh medium. To obtain an enriched population of titan-like cells, we elutriated the cultures as described in M&M to recover the cells of larger size. After isolation of the RNA, we investigated gene expression by RNAseq. After mapping the reads and subsequent analysis, we identified genes that were overexpressed or repressed in titan-like cells by two-fold in the different replicas performed (Fig 11A and supplemental table 2). Interestingly, the gene that was most overexpressed in titan-like cells was CNAG_01653. This gene encodes the mannoprotein Cig1 (CNAG_01653, initially annotated as cytokine inducing glycoprotein), which is involved in iron uptake from heme groups and is overexpressed during iron limitation conditions [46, 47]. We also found that the transcription factor Rim101, (which is required for titan cell formation and is activated by PKA [27, 48]) was also upregulated in titan-like cells. Among the repressed genes, we found *PCL1*, which encodes a cyclin that regulates the transition between G1 and S phases in the cell cycle. There were also some genes that were repressed in titan-like cells involved cell wall rearrangements, such as chitin deacetylases.

**Figure 11:**
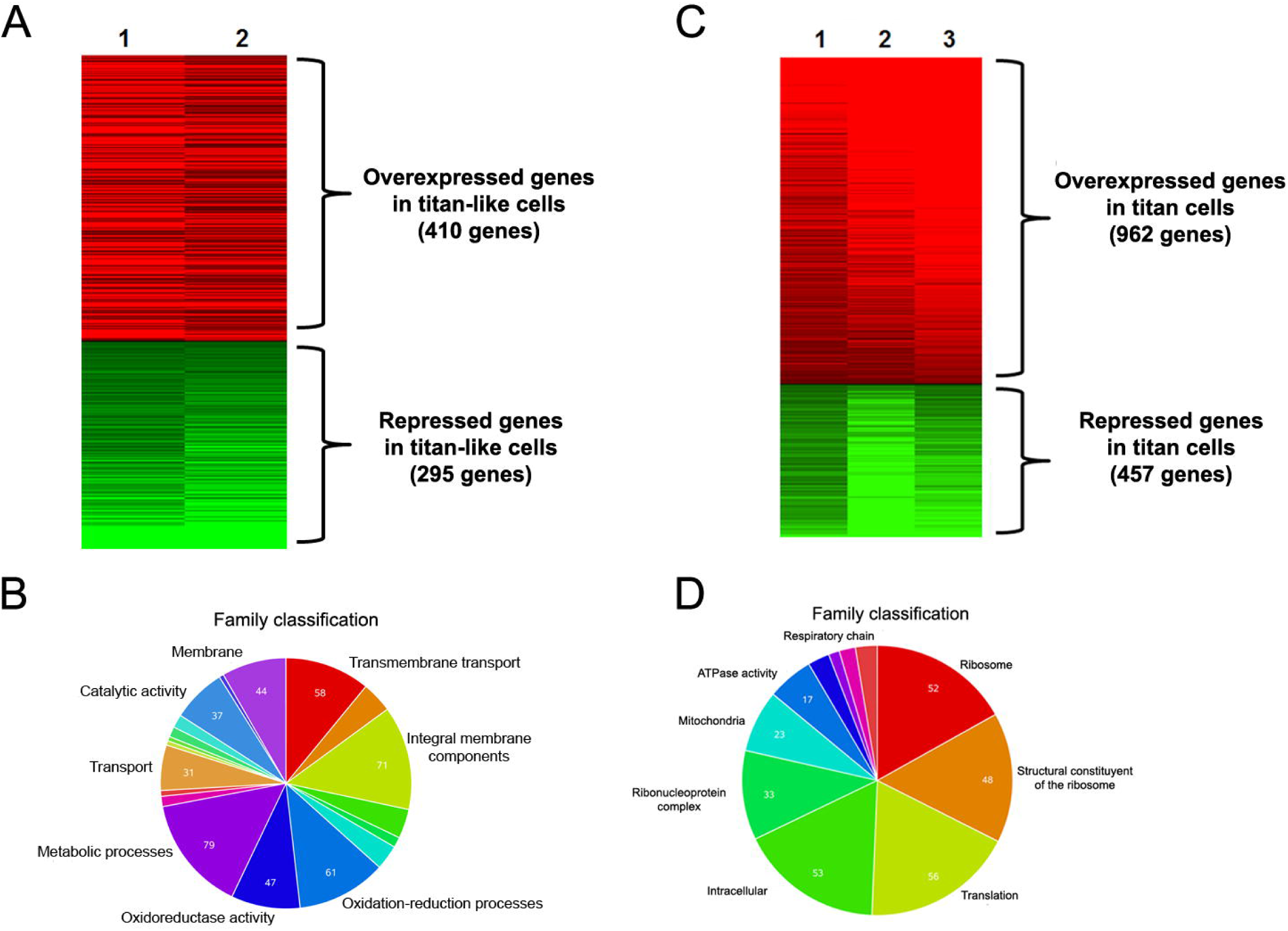
Representation of gene expression changes in RNAseq experiments in titan-like cells obtained in vitro and titan cells isolated from lungs. A-B) Analysis of gene expression changes of titan-like cells obtained in vitro. Titan-like cells were obtained in TCM with shaking and transferred to new medium every 24 h for three days. Parallel cultures were carried out in Sabouraud. The cultures were incubated at 37 °C, and after this time, titan-like cells were isolated by elutriation as described in material and methods. RNA was isolated and RNAseq was performed and analysed as described in M&M. Data from two biological replicates was obtained. A) Heat map representation of the reproducibility of the fold change in the different replicas. Color map at the left represents the fold-change represented by each color (log_2_ scale). B) Analysis of overrepresented families among the genes that were significantly overexpressed by at least two fold in all the replicas. The graph was obtained using FungiFun webpage. C-D) Analysis of gene expression changes of titan cells isolated from the lung. Twenty C57BL/6J mice were infected with H99 strain, and after 14 days, cryptococcal cells were isolated from the lung as described in material and methods. Titan cells and small cells were separated by elutriation and RNAseq was performed. C and D represent the same analysis described in A and B. The infections and RNA isolations were performed in three experimental sets.

We performed an in silico functional analysis of the genes that were overexpressed (>2 fold) in titan-like cells using FungiFun (see M&M). In this analysis, we found a large number of membrane protein and receptors (Fig 11B).

One limitation for obtaining titan-like cells *in vitro* is that their size does not become the same as titan cells that form in the lungs. For this reason, we decided to investigate the gene expression changes in titan cells obtained from lungs of infected mice. For this experiment, we infected C53BL/6J mice because we previously demonstrated that in this genetic background there is a higher proportion of titan cells compared to other mouse strains [23]. After two weeks of infection, we sacrificed the animals and isolated the yeast population as described in M&M. Since the cryptococcal size *in vivo* is very heterogeneous, we separated titan cells from cells of regular size by elutriation, so in this way we could compare gene expression of the two populations exposed to the same environmental conditions (lung). After RNA isolation, we performed again RNAseq using Illumina Technology. When we examined the results, we unfortunately found that the number of reads that mapped in the *C. neoformans* genome was low (around 2x10^5^ to 2x10^6^, depending of the sample), which was a limitation to perform a deep gene expression study. We identified the genes that were upregulated or repressed by two-fold (Fig 11C and supplemental table 3). We found that the most expressed gene with known function in titan cells encodes an extracellular elastinolytic metalloproteinase. In addition, many genes encoding proteins related to mitochondrial activity were found, as well as the iron metabolism-related encoding gene *CIG1* mentioned above. As we did in the analysis of titan cells in vitro, we performed a functional analysis of the genes that were overrepresented among overexpressed genes. As shown in Fig 11D, in this case, we found a large number of genes encoding proteins with metabolic functions, in particular, respiration and protein synthesis.

We finally compared the genes that were overexpressed or repressed in both titan-like cells obtained in vitro and in titan cells isolated from the lungs. We found that there were 80 genes that were upregulated in both types of cells, and 21 that were consistently repressed. Among the genes that were overexpressed in titan-like and titan cells, we found mainly metabolic genes (in particular, related to triglyceride metabolism) and some cell wall remodeling enzymes (such as 1,4-α-glucan-branching enzyme). In addition to CIG1, we also found other genes related to iron metabolism, such as a high affinity iron transporter (*FTR1*).

## DISCUSSION

*Cryptococcus neoformans* is an exceptional model to understand mechanisms induced by pathogenic fungi to adapt to the host and cause disease. In this sense, this yeast has developed specialized phenotypes that confer a clear advantage in the lung environment. Some of the most important changes are related to changes in cell size, which can occur by growth of the capsule, or growth of both the cell body and capsule. In this last case, *C. neoformans* induces a specific cell type that has been denominated as titan cells [20, 21], although their function in the virulence and the mechanisms that regulate their formation are not known. At the moment, only the cAMP-dependent and the mating pathways have been described as relevant in titan cell formation [20, 21, 27]. In this sense, it has been shown that increased expression of the PKA induces cell enlargement in *C. neoformans* and the ploidy of the cells [49, 50]. One of the limitations to study titan cells is the difficulty of reproducing this phenomenon *in vitro*. Originally, we described that a small proportion of titan cells could be observed in minimal media [21]. Then, it was described that these cells appear in the presence of phospholipids [28]. In this work, we have defined a culture medium in which we have replicated this phenomenon in vitro. This medium has several characteristics: limitation of nutrients at neutral pH, inclusion of mammalian serum and a CO_2_ enriched atmosphere. In these conditions, we could obtain a high proportion of cells of a size around 30 μm (capsule included). This diameter is smaller than the average size measured in mouse lungs [11, 20, 22], where average size is around 40-50 μm, but that can reach up to 100 μm, so we argue that in our TCM we could be examining the initial steps of this morphological transition. It is possible that *in vitro* the cells do not reach the same size than cells obtained from *in vivo* infections because the nutrients of the medium or the factors that favor this morphological change are consumed. Instead, *in vivo* infections are maintained for a longer time, with a stable nutrient concentration allowing cells to reach significantly larger sizes. Despite this limitation, the availability of *in vitro* conditions that, at least in part mimic titan cell formation is a key contribution to understand the biology of these cells.

In general, our results indicate that titan-like cell formation is induced by multiple factors, being some of them necessary, but not sufficient, to trigger the transition. This is the case of serum because it only induced cell growth under nutrient limiting conditions, but not in rich media. Similarly, titan-like cells were not observed after incubation with 100% serum. These results indicate that titan-like cells are formed in response to some elements of the host present in the serum in the context of the stress produced by the limitation of nutrients in the medium. These findings allow the dissection of the intracellular pathways that are triggered during cellular growth.

It has been described that serum and the fraction of polar lipids induce cell enlargement in *C. neoformans*. Our results agree with previous findings [28], because polar lipids were also able to induce the formation of titan cells in TCM. The most plausible mechanism for this mechanism is that phospholipids are degraded by phospholipase C, which produces diacylglycerol (DG), or by phospholipase B, which produces arachidonic acid (AA) [51]. DG activates human as well as *C. neoformans* PKC [34]. To investigate the role of this signaling pathway, we blocked the activity of this kinase with several pharmacological inhibitors, and found that both staurosporine and calphostine C had a dramatic effect on titan-like cell formation. This protein participates in multiples processes, such as maintenance of the integrity of the cell wall [52-54] and polarized growth [55]. Mutants lacking the *PKC1* gene present many cellular alterations, such as osmotic instability and susceptibility to temperature [35]. We tried to evaluate the effect of these inhibitors at a lower temperature and in the presence of sorbitol in an attempt to overcome the cellular defects associated with the absence of Pkc1. However, we found that sorbitol inhibited titan-like cell formation (results not shown), suggesting that osmotic pressure impairs cell growth. For this reason, further studies are required to understand how this pathway participates in titan-like cell formation.

Nutrient limitation is required for the serum-induced development of titan-like cells, which indicates that titan-like cell induction is a response to stress factors. Both the nutrient concentration in the medium and the carbon source induce multiple changes in the cell. For example, high concentrations of glucose induce catabolic repression of genes required for the use of other carbon sources [56, 57]. Furthermore, the activity of the TOR pathway (*Target of Rapamycin*) is regulated by nutrient availability [58]. TOR proteins are highly conserved kinases that regulate cell growth, cellular division and respond to changes in the environment such as nutrient availability or cellular energy status [59]. In recent years, it has been demonstrated that the TOR route controls different processes, all related to cell growth, membrane trafficking, protein degradation and signaling through PKC [60]. Our data suggests that this morphological transition requires TOR signaling pathway. It has been shown that these kinases are active in the presence of nutrients and stimulate growth, although the exact role of this pathway in *C. neoformans* is not known. In fact, it would be expected that inhibition of TOR would inhibit cell replication and increase cell size due to the cell cycle arrest. We believe that these results support the idea that titan-like cells are formed by endoreduplication, so several rounds of cell cycle progression associated with absence of M phase are required for cell growth. Moreover, it has been shown that the TOR pathway positively regulates PKC phosphorylation [61], so a decrease in activity of TOR proteins might influence titan-like cell formation through a defective activation of PKC.

We also observed that other important factor for titan-like cells formation *in vitro* is CO_2_. The capsule is induced in response to CO_2_ concentrations present in the host [62], which is consistent with this factor also favoring the growth of the cell body. Carbonic anhydrase convert CO_2_ into HCO_3_^-^, which activates adenylate cyclase [44, 63], so our results are consistent with the hypothesis that CO_2_ induces cell growth through the cAMP pathway. In *C. neoformans*, there are two carbonic anhydrases encoding genes, *CAN1* and *CAN2*, being Can2 the most important [44]. Interestingly, deletion of *CAN1* resulted in an enhancement of the production of titan-like cells. Although we do not know the molecular mechanism for this phenomenon, we postulate that in the absence of Can1, there might be a compensatory overexpression of Can2 that could induce a stronger activation of the cAMP pathway.

We also observed that subinhibitory concentrations of azide have a modest, but reproducible positive effect on titan-like cell development. We argue that a partial inhibition of the respiratory chain can trigger a stress signal that results in a stop of the cell cycle and allows cellular size increase. In this regard, it could be assumed that a limitation in the respiratory capacity might lead the organism to generate energy, at least in part, by fermentative metabolism, a situation that has been associated to increased PKA activity in many fungi [64]. Although *C. neoformans* is mainly a respiratory yeast, it has been shown that it can produce both ethanol and acetate in vitro and in vivo [65, 66], so a similar activation of the PKA activity could also occur in these conditions.

One of the most striking results of this work is the effect of cell density on the formation of titan-like cells, suggesting that this transition is regulated by *quorum sensing* (QS) phenomena, a system of cell-cell communication mediated by molecules that are released into the medium by the cells. In this way, a higher cell density results in a higher concentration of released molecules and greater effect on the cells [29, 67, 68]. Farnesol was described as the first molecule responsible for QS phenomena [69]. In *C. albicans*, farnesol represses the transition from yeast to hyphae [69]. In *C. neoformans*, this molecule inhibits growth *in vitro* and has an effect on the secretion of phospholipases and proteases [70]. Previous reports demonstrate that QS regulates the formation of biofilms, the release of the capsular polysaccharide glucuronoxylomannan (GXM) and the melanin synthesis of this pathogen [30]. Under our conditions, farnesol inhibited the formation of titan-like cells. This molecule has been reported to inhibit adenylate cyclase in *C. albicans* [71] so we believe that the effect of this compound could be mediated through a decrease in cAMP concentrations in the cell.

A major QS signal for *C. neoformans* is a 11-mer peptide (Qsp1) [31, 32]. We tested the effect of this peptide on titan-like cell formation, and found that it inhibited the transition. Qsp1 promotes cell division and replication. Although further studies are required to understand the molecular mechanism by which Qsp1 regulates this transition it is plausible to propose that, since titan-like cell are formed in the absence of budding, Qsp1 blocks this development due to its positive effect of cell division and replication. The biological role of QS phenomena on titan-like cells is not known, Furthermore, some morphological transitions in other species, such as hypha development is *Candida albicans*, are repressed by QS molecules. It is worth noting that a cell-dose relationship similar to that found here has been described *in vivo*, since a low number of cryptococcal CFUs in the lung correlates with a higher proportion of titan cells [21]. Therefore, it will be necessary to elucidate the role of Qsp1 in the context of lung colonization in the future. However, this peptide does not seem to be the key factor that represses cellular growth at high densities, since *qsp1* mutants form titan-like cells similarly to the wild type strain. In agreement, Albuquerque et al described that *C. neoformans* is able to produce QS molecules that are not susceptible to high temperature, proteinase, trypsin, pronase, DNAse, RNAse and glucosidase [30]. These authors also described that farnesol, tyrosol and Qsp1 did not replicate the effects observed with their conditioned media, and demonstrated that pantothenic acid could in part reproduce QS phenomena. In summary, we hypothesize that multiple QS molecules (farnesol, Qsp1 and others), negatively regulate titan-like cell formation. The fact that some of these molecules are not produced by *C. neoformans* suggests that QS phenomena induced by other microorganisms of the environment and natural flora could interfere with the adaptation of *C. neoformans* to stress conditions.

In summary, we have found several conditions that promote cellular enlargement in *C. neoformans*. In Fig 12, we present a suggestive and simple representation on how the different factors and pathways described in this work could exert their action and how they could regulate titan-like cell formation.

**Figure 12:**
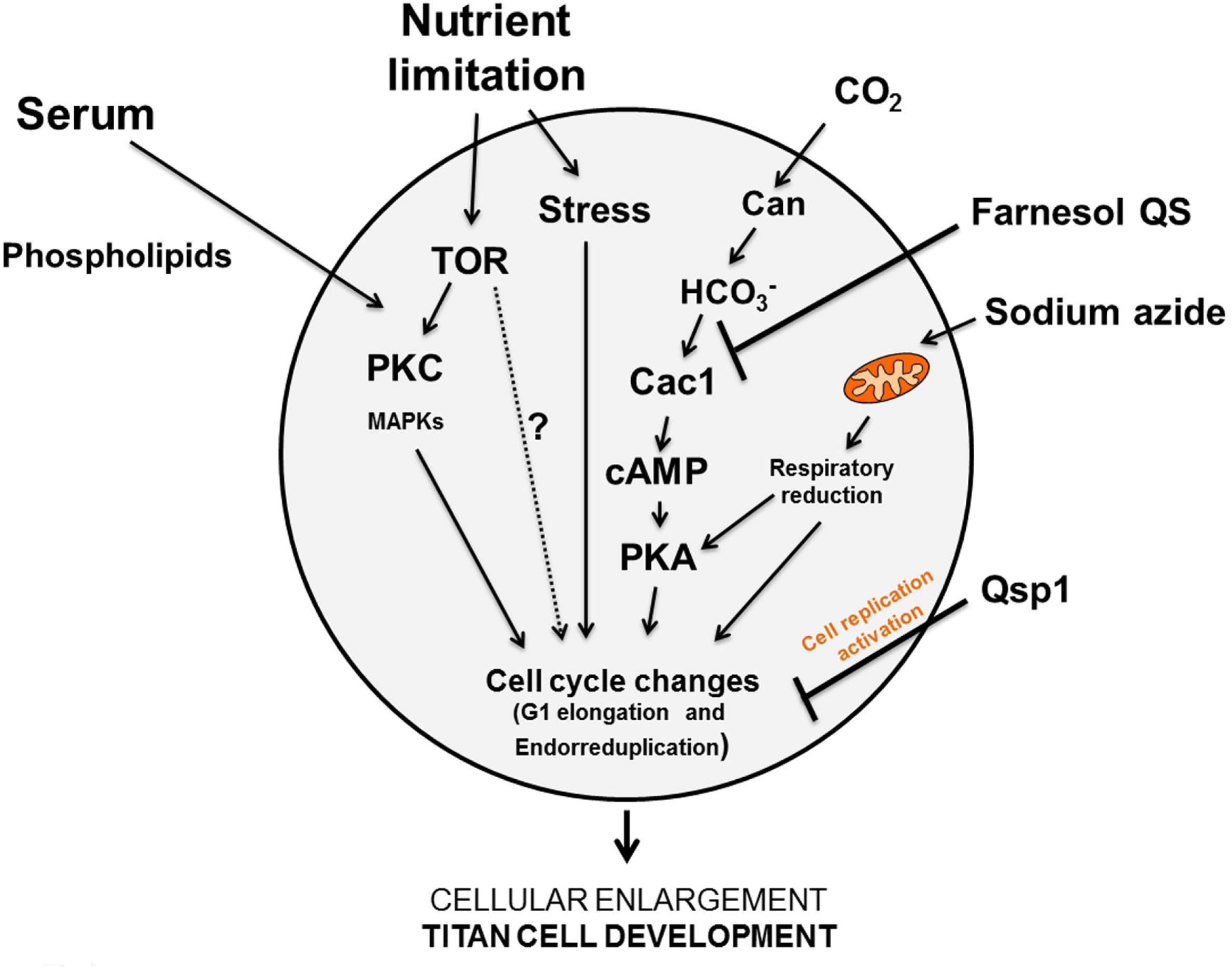
Schematic representation of the integration of different factors of the TCM medium affecting titan-like cell formation and the intracellular pathways that could be activated or repressed.

The formation of titan cells has been observed in clinical samples [72-74], and despite being a mechanism that confers advantages to *C. neoformans* against the host during infection, we observed that this process is not a universal phenotype. Our results showed that there is great variability among different isolates, and not all the strains did form titan-like cells in vitro. Our work provides a new way to investigate the genetic differences between strains with high and low capacity to form titan-like cells using genomic approaches. However, the strains of *C. neoformans* var. *grubii* (serotype A) are those where the proportion of titan-like cells was higher, suggesting that this serotype has a greater capacity of adaptation to the lung. This result is in agreement with the literature, since this serotype is isolated more frequently in infected patients [10], suggesting that there is a correlation between the ability to form titan cells and development of the disease. We believe that this correlation should be confirmed in future clinical studies.

An interesting finding of our work is the impaired ability of acapsular mutants or strains affected in capsule enlargement to form titan-like cells. This finding supports the idea the capsule synthesis is regulated by factors that also affect cell size. Previous reports demonstrate that after capsule growth, the size of this structure correlates with cell body size [75]. In addition, capsule growth mainly occurs in G1 [76], which is also the cell cycle phase in which the growth of the cell body occurs. All these data indicate that capsule growth and cell size are linked. The fact that the alteration in capsule synthesis affects cellular growth also suggests that these two processes share common pathways.

Titan cells contribute to virulence through different mechanisms, such as polarization of Th2-type immune responses [23, 24], replication [26, 77], resistance to oxidative damage [20, 21] and phagocytosis avoidance [21, 25]. In our case, titan-like cells obtained in vitro were not phagocytosed, most probably due to their large size. In our case, these cells did not prevent the phagocytosis of cryptococcal cells of regular size, as it is the case in experiments performed *in vivo* [25]. This difference can be explained by multiple factors. We believe that activation of the macrophages *in vivo* is different from *in vitro*, so in the lungs there could be multiple factors that affect the phagocytic activity of macrophages, explaining why the effect of titan cells *in vivo* on macrophages are not identical from the effects induced *in vitro*. Furthermore, it is also possible that titan cells *in vivo* express virulence factors or epitopes that are not produced in vitro, so they may have a different effect on macrophages. Further studies are required to fully understand the response of immune cells to cryptococcal titan-like cells.

The isolation of titan-like cells *in vitro* opens new perspectives and research lines. For example, we investigated gene expression changes associated with titan cells development. Our approach had, however, some limitations. Since titan-like cells were mainly obtained in static conditions in cultures incubated in a CO_2_ chamber at low cell densities, it was difficult to isolate enough RNA to perform RNAseq. So we decided to obtain titan-like cells from TCM but with shaking after passaging to fresh medium during several days. In this way, we found that titan-like cells overexpressed the gene that encodes Cig1, which is a mannoprotein involved in iron uptake from heme groups. These findings suggest that iron concentration in the medium can also regulate the induction of titan-like cells. Interestingly, iron limitation also induces the growth of the cryptococcal capsule [78]. *CIG1* expression is activated by the cAMP-regulated transcription factor Rim101 [46], which was also upregulated in titan-like cells. This transcription factor is required for titan cell formation in vivo [27]. In consequence, our transcriptomic data is agreement with a key role of the cAMP pathway in titan-like cells, which was supported by the fact that *cac1* mutants do not form this type of cells in vitro (figure 8B). In addition to *CIG1*, we found a significant overexpression of other membrane proteins and transporters encoding genes, suggesting that titan-like cells induce a response to capture nutrients and solutes from the medium to obtain enough energy to enlarge cell size.

Titan cell formation occurs by endoreduplication, so presumably all the factors and signaling pathways must alter the normal progression of cell cycle. Interestingly, we found that in titan-like cells, the G1/S cyclin Plc1 was repressed, which suggests that the process requires an elongation of the G1 phase in addition of the absence of mitosis. This result is in agreement with previous findings that demonstrated that *plc1* mutants have increased capacity to form titan-like cells *in vivo* [27]. In our titan-like cells, we found that they have a higher content of DNA compared to cells of regular size, suggesting that during their development requires endorreduplication events as occurs *in vivo*. Our imaging of the process (supplemental video 2 and 3) also suggests that this cell cycle alteration is accompanied by an elongation of the G1 phase, which is the cell cycle phase in which the cell mainly enlarge their size. We believe that our data provide a new approach to characterize in future studies the cell cycle alterations that trigger titan cell development.

Since titan-like cells obtained *in vitro* did not reach the size of the titan cells found in the lungs, we tried to investigate the gene expression profile in these cells. Our approach has the advantage that we are comparing changes in gene expression between cells of normal cells and titan cells isolated from the same host environment (lung). However, due to the complexity of the protocol, we also are aware that our results have the limitation that the number of reads obtained was low to perform a deep analysis. Still, it was noticeable that the genes mostly expressed in titan cells encoded an elastinolytic metalloprotease that is secreted to the medium [79]. Extracellular cryptococcal proteases have been involved in multiple processes required for virulence, such as degradation of host extracellular matrix [80] and crossing of the BBB and brain invasion [81, 82]. Although presumably titan cells do not disseminate to the brain due to their size, the fact that they overexpressed this metalloprotease *in vivo* is interesting because it suggests that they can modulate the host extracellular medium to favor their survival in the lung. The expression of this gene has been shown to be controlled by Gat201 during growth in DMEM [83], which is also required for titan-like cell formation (see Fig 8A).

We also observed that *in vivo* titan cells overexpressed genes that encode metabolic and mitochondrial proteins, which could reflect changes in metabolism required for proper energy production required for cellular enlargement. Many of these genes were related to Acetyl-CoA metabolism, which has been shown to be important during metabolic adaptation of C. neoformans during infection [84]. Interestingly, we also found several genes related to iron metabolism (*CIG1* and *FTR1*), so further studies are warranted to investigate the role of this metal on titan cell formation.

Finally, we would like to acknowledge that other groups (Dr. Alanio, Pasteur Institute, France, and Dr. Ballou, University of Birmingham, UK) have identified in parallel to our work other conditions in which titan-like cells are formed, mainly based on minimal media. We believe that these works together with our findings provide significant advances to the scientific community because they will allow the design of multiple research lines that will facilitate the characterization of the factors and signaling pathways that are involved in titan cell development. The data presented by different groups also indicate that *C. neoformans* may induce titan-like cells in vitro in response to multiple factors. In addition, the ability to obtain titan cells *in vitro* will also have a positive bioethical impact because we will be able to significantly reduce the number of experimentation animals that it was previously required to characterize these cells.

## METHODS AND MATERIALS

### Strains and media

In most of the experiments, we used *C. neoformans* var. *grubii* (serotype A) H99 strain [85], but we also included, *C. neoformans* var. *neoformans* (*C. deneoformans*, serotype D), A/D hybrids and *C. gatti,* and different mutants obtained from the library described by Liu and coworkers [41] and from the Fungal Genetic Stock Centre. All strains are described in table 1. Strains were preserved in Sabouraud medium containing 30% at -80% and were recovered at 30 °C in Sabouraud solid medium (Oxoid LTD, UK).

**Table 1.**
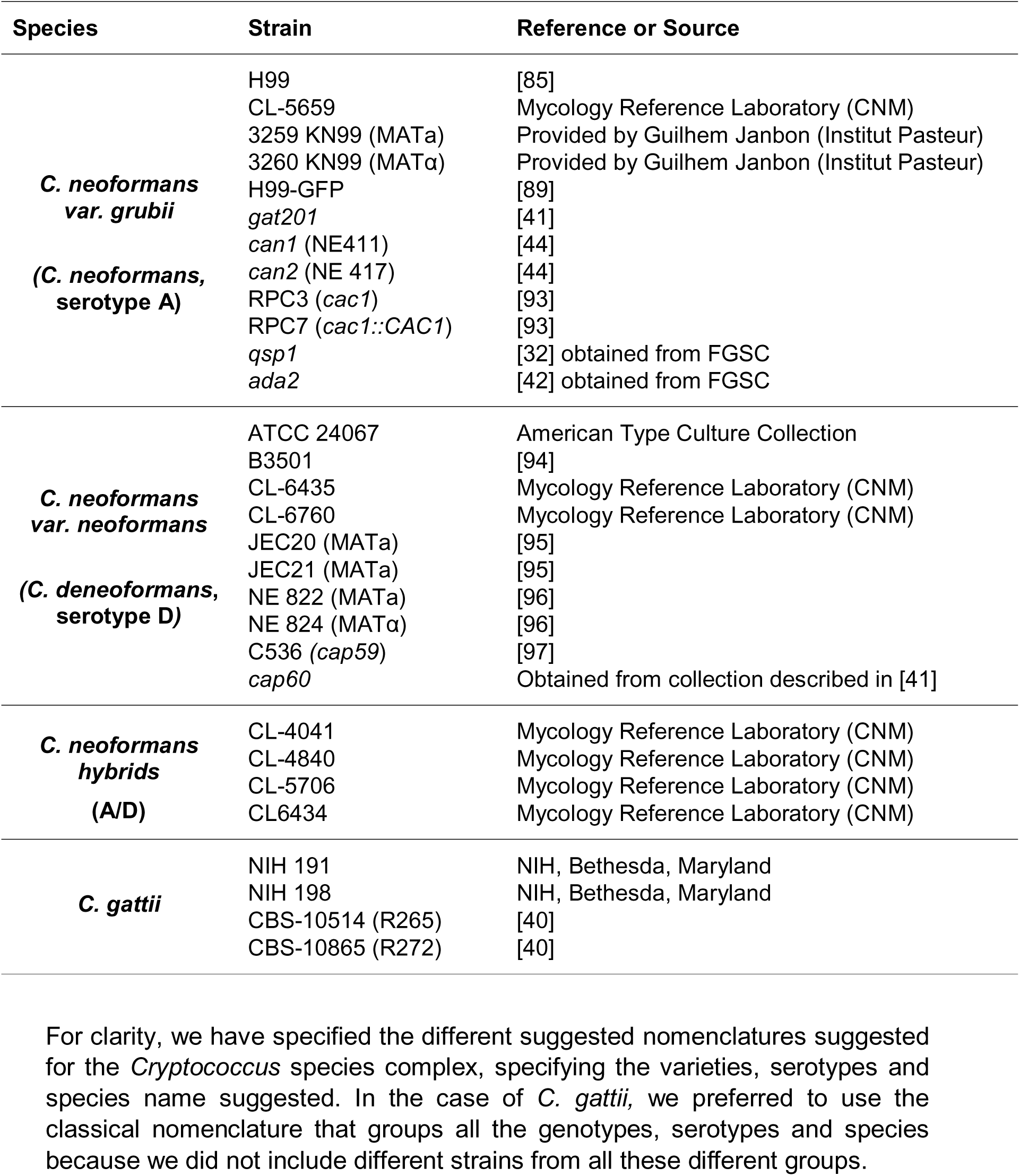
List of strains used

The yeasts were routinely grown in liquid Sabouraud medium at 30 °C or 37 °C with shaking (150 r.p.m.). To induce titan-like cells, strains were grown in a medium that we have defined as Titan Cells Medium (TCM), which is based in the medium described to induce capsule growth (10% Sabouraud buffered at pH 7.3 with 50 mM MOPS, [86]). TCM contains 5% Sabouraud and 5% inactivated fetal calf serum (FCS, Biological Industries) diluted in MOPS 50 mM at pH 7.3 plus 15 μM sodium azide (Sigma Aldrich). Cultures were grown in tissue culture flasks or 96-wells plates at 37 °C in an atmosphere enriched with CO_2_ for 18 hours.

### India Ink staining and measurement of cell size

To observe and measure the size of the cells, 10 μL of a cell suspension were mixed with a drop of India Ink drop (Remel Bactidrop, Lenexa, Kansas) and observed in a Leica DMI 3000B microscope. Pictures were taken with a Leica DFC 300FX camera using the Leica Application Suite (Leica Microsystems) and processed with Adobe Photoshop 7.0 (San Jose, CA). [86].

### Lung infections of mice with *C. neoformans*

Six to eight weeks-old male mice from C57BL/6J (in house bred at the National Centre for Microbiology) were used in all experiments. The animals were kept in ventilated racks at 22-24°C with proper environmental enrichment (cupboard houses and hollow cylinders).

Yeast cells were incubated overnight in liquid Sabouraud medium at 30 °C, centrifuged at 2830 g, washed and suspended in sterile PBS. The cell density was determined using a TC20 cell counter (BioRad) and a suspension of 3.3x10^7^ cells/mL was prepared in sterile PBS. The animals were anesthetized with a mixture of ketamine (Imalgene 1000, 50 mg/Kg) and xylazine (Xilagesic 2%, 20 mg/Kg) and infected intranasally with 30 μL of the yeast suspension (10^6^ cells per mouse) as previously described [23]. Animals were sacrificed after 14 days of infection by exposure to a high CO_2_ enriched environment.

### Organ extraction

In all experiments, we performed a perfusion to remove blood from the lungs. A cut was made in the jugular vein of the mice and 5-10 mL of PBS were injected in the left ventricle of the heart. Then, we excised and homogenized the lungs in 10 mL of sterile water using cell strainers (100 μm size pore, BD Falcon) and a 5 mL syringe plunge in Petri plates. This process disrupts the mammalian cells without significantly affecting the integrity of the cryptococcal cells. Then the cell suspension was centrifuged at 2,830 g and washed with sterile water three times to fully break and remove the mouse cells. Finally, the yeasts were suspended in PBS.

### Cell separation by elutriation

Titan and normal cells from *in vivo* assays were separated using an Avanti® J Series instrument from Beckman Coulter with a JE-5.0 elutriator rotor. Counter-current centrifugal elutriation is a process for separating a heterogeneous cell suspension into its constituent cell populations according to their size, shape and density [87]. Briefly, the cellular suspension obtained from *in vivo* experiments as described above was introduced into the elutriation chamber at 4 °C. Cells were recovered in different fractions of 100 mL, by increasing the flow rate, and fractions were kept on ice. A total of 11 fractions were collected and centrifuged at 2830 g. The efficiency of elutriation was confirmed first by determination of the total size of the cells with India ink and second, by flow cytometry. The first 4-5 fractions mainly contained cells of normal size (4-7 μm), where fractions 8-11 contained cells with a size above 30 μm (titan cells). For this reason, fractions 1-4 and 8-11 were pooled (small cell fraction and titan cell fraction respectively). The elutriation conditions are described in supplemental table 1.

### Growth curves

Yeast strains were grown in 10 mL Sabouraud liquid medium with 10 μL of 15 mM sodium azide. The cultures were grown at 30 °C with continuous shaking (150 r.p.m) overnight. Cells were harvested with centrifugation at 2,830 g for 3 minutes and washed twice with PBS. Cellular suspensions were prepared at 2x10^5^ cells/mL in Sabouraud or TCM using a TC20 Automated Cell Counter (BioRad). One hundred and seventy μL of the cultures were placed in a 96 well plate in triplicate and were incubated at 30 °C or 37 Since titan-like cells obtained C during 24 hours without shaking in a spectrophotometer iEMS (Thermofisher). Optical density was measured at 540 nm every hour for 48 hours. Data were analyzed with Graph Pad Prism 6.

### Lipids extraction of Fetal Bovine Serum

Polar lipids from serum were obtained as described in [28]. Briefly, aliquots of 1 mL of Fetal Bovine Serum (FBS) were shaken with a mixture of chloroform and methanol (2:1) (v:v) for 3 hours at room temperature. The samples were centrifuged for 10 minutes at 2,500 r.p.m for phase partitioning. The upper phase was collected in a 1.5 mL tube and dried during 1 hour in a SpeedVac concentrator. The pellet was suspended in 200 μL of PBS and conserved at 4 °C. To evaluate the effect of fetal calf serum polar lipids on titan-like cell formation, we performed experiments with different amounts of the extraction solution described above (1/40; 1/100 and 1/200 dilution) in 5 % Sabouraud buffered at pH 7.3 with 50 mM MOPS and 15 μM sodium azide. As a control, the same medium with PBS was used. In parallel, cells were grown in Sabouraud and TCM as growth control and titan-like cell formation respectively.

### Visualization of titan-like cell formation by real-time microscopy

Yeast cells were inoculated in TCM at 10^4^ cells/mL as detailed above. One hundred seventy μL from the yeast suspension were placed in a 96 well plate and incubated at 37°C with 5% CO_2_ under a Leica DMI 4000B microscope. Photographs were taken every 3 min using the 20x objective. The videos generated by the Leica software were exported as AVI documents and processed with ImageJ software. In all cases, the videos were assembled with 12 frames per second, so one second of video corresponds to 36 minutes of real time. Time was included in each frame was included using the Time Stamper plugin from ImageJ.

### Quorum sensing assays

Cell suspensions were prepared at a concentration 10^6^ cells/mL in parallel in Sabouraud with 15 μM sodium azide and TCM. Serial 1/10 dilutions were made up to 10^3^ cells/mL. A volume of 170 μL of these suspensions was incubated in a 96 well plate at 37 °C overnight with 5% CO_2_ without shaking. The plates were observed in a Leica DMI 3000B microscope. Pictures were taken with a Leica DFC 300FX camera using Leica Mycrosystems software. The cell body diameter of thirty to fifty cells was measured with Adobe Photoshop 7.0.

### Effect of conditioned media on titan-like cell formation

*Cryptococcus neoformans* was inoculated at 10^4^ and 10^6^ cells/mL in TCM as described above and incubated at 37 °C in a 5% CO_2_ enriched atmosphere. After 18 h of incubation, the cultures were centrifuged and the supernatants collected to yield Titan-like Cells Supernatant (TCS) and regular cells supernatant (RCS). To evaluate the influence of these supernatants on the titan-like cell formation, *C. neoformans* cultures inoculated at 10^4^ cells/mL in 96 wells plates were prepared in different growth conditions: 1) Fresh TCM medium (TCM), 2) Supernatant from cultures of titan-like cells (TCS), 3) Supernatant from cultures of cells of regular size (RCS). These conditioned media were mixed with fresh TCM (1:1 proportion v/v). As control, we carried out a culture in which fresh TCM was diluted with the same volume of distilled sterile H_2_O. After the inoculation of the different media and mixtures with *C. neoformans* at 10^4^ and 10^6^ cells/mL and incubation at 37 °C in a CO_2_ incubator for 18 h, the cell size was measured by microscopy as described above.

### Peptide chemical synthesis

Chemical synthesis of the peptides was done by the proteomic facility of the National Centre for Biotechnology (CSIC, Spain) using an Multipep automatic synthesizer (Intavis, Köln, Germany) and Fmoc-Amino Acid Wang resins (Merck, Darmstadt, Germany). After release from the resin, the peptides were purified by reverse-phase chromatography in a semipreparative HPLC system (Jasco, Tokio, Japan) with a C18 Kromaphase column (Scharlab, Barcelona, Spain). The fractions were analyzed by mass spectrometry and lyophilized until their use. We synthesized peptides described in [32]: Qsp1 (NFGAPG**G**AYPW), an inactive version of this peptide (NFGAPG**A**AYPW) and a scrambled Qsp1 peptide (AWAGYFPGPNG). The peptides were dissolved in sterile PBS at 1 mM, and their effect on titan-like cell formation was tested at 30 μM and 15 μM in TCM. The samples were incubated for 18 hours at 37 °C with 5% of CO_2_ in 96 wells plates. After the incubation period, the cells were observed by optical microscopy and the body cell sizes were measured.

### Effect of Farnesol on titan-like cell formation

To test the effect of Farnesol on titan-like cell formation, we followed the protocol described in [70] with some modifications. Briefly, we prepared Farnesol at a concentration of 400 mM in 7.5% of dimethyl sulfoxide (DMSO, Sigma-Aldrich). Then, we prepared the Farnesol stocks (50x) at an initial concentration of 15.04 mM. Afterwards, we made 1:2 serial dilutions of Farnesol in DMS0 at 3.75% to a final concentration of 0.029 mM. The yeasts were grown in Sabouraud+azide overnight at 30 °C and the inocula were prepared at 10^4^ and 10^6^ cells/mL in Sabouraud+azide and TCM. Dilutions (1:50) from each stock concentration of Farnesol were made in 96 well plates (Costar, New York) containing the yeast in Sabouraud and TCM (final volume of 170 μL). The plates were incubated at 37 °C overnight with 5% CO_2_. In this way, the final concentrations range of Farnesol used was from 300 μM to 0.5 μM, leaving the concentration of the solvent (DMSO) at 0.075%, which we tested that did not interfere with titan-like cell formation. The plates were observed in a Leica DMI 3000B microscope and the optical density was measured at 540 nm at 24 and 48 hours using a spectrophotometer iEMS (Thermofisher).

### Nuclei analysis in titan-like cells

*Cryptococcus neoformans* cells from H99 strain were cultured overnight in TCM at 10^4^ and 10^6^ cells/mL at 37 °C with CO_2_ without shaking. After the confirmation of titan-like cells formations by optical microscopy, cells were washed with dH_2_O and fixed with 70% ethanol at 24 °C for one our followed by an overnight incubation at 4 °C. After fixing, the cells were washed twice with RNAse A buffer (0.2 M Tris, pH 7.5, 20 mM EDTA) and treated with 10 μg/mL of RNAse A for 4 hours at 37 °C. After the incubation, cells were washed twice with PBS, suspended in PBS and incubated overnight at 4 °C. Next day, the cells were centrifuged and suspended in a 200 ng/mL solution of 4’,6-diamidino-2-phenylindole (DAPI) and incubated in the dark for 10 minutes at room temperature to stain the nucleus. Then, the cells were washed, suspended in PBS and the fluorescence intensity of the nucleus were analyzed by flow cytometry. Cells were examined for cell size by forward scatter parameter (FCS) and granularity by side scatter parameter (SSC) using the BD LSRFortessa X-20cytometer (BD, Bioscience). Two populations of titan-like and regular cells were delimited and, in each population, the fluorescence intensity of the DAPI staining in 10,000 cells was measured. Data obtained were analyzed with the BD FACSDiva (BD, Bioscience) and FlowJo 7.6.1 (Tree Star Inc, Ashland, Oregon) softwares. The nucleus of the *C. neoformans* H99 titan-like and regular cells stained with DAPI were also observed by confocal microscopy using a Leica SP5 confocal microscope.

### Influence of the protein kinase C (PKC) inhibitors and rapamycin on the formation of titan-like cells

The influence of the PKC pathway in the formation of the titan-like cells was evaluated by the addition of four different inhibitors: calphostin C, staurosporine and bisindolylmaleimide I, and genistein (all from Calbiochem) that inhibits tyrosine kinase as a control. For this purpose, 10 μM, 5 μM and 1 μM of calphostin C, bisindolylmaleimide I and Genistein and 0.01 μM and 0.001 μM of staurosporine. In all cases, the final concentration of DMSO was 0.1%, which did not inhibit titan-like cell formation. To inhibit TOR kinases, we used rapamycin at a final concentration of 1 μM. All the inhibitors were prepared in DMSO, so parallel controls with the same concentration of solvent (0.04%) were carried out.

### Phagocytosis of titan-like cells *in vitro* with RAW 264.7 macrophages

RAW 264.7 macrophages were maintained in DMEM medium supplemented with heat-inactivated 10% fetal bovine serum (FBS, Hyclone-Perbi), 10% NCTC medium and 1% non-essential amino acids (Sigma-Aldrich, Steinheim, Germany). The day before the experiment, the macrophage monolayer was separated from the plate by pipetting and the cells were centrifuged at 1,265 g. Macrophage suspensions were prepared at 2.5x10^5^ cells/mL. Two hundred μL per well were inoculated into 96 well plates and incubated overnight at 37° C and 5% CO_2_. The next day, different types of cells (titan-like cells obtained *in vitro*, and cells of regular size) at a final concentration of 5x10^5^ cells/mL with 5 μg/mL of monoclonal antibody 18B7 [88] were added to the macrophages for 2 hours. Phagocytosis was quantified by two different methods. First, the plates were observed with a Leica DMI 3000B microscope, and the percentage of infected macrophages was determined visually. In some cases, the plate was visualized with a Leica 4000B with a chamber that allowed incubation at 37 °C and 5% CO_2_, and videos were taken as explained above. Alternatively, we quantified the phagocytosis percentage by flow cytometry. In this approach, we performed phagocytosis assays as above, but using larger volumes in 24-well plates containing 1 mL of medium. To differentiate yeast cells, we used a H99 strain that expresses the green fluorescence protein (H99-GFP) [89]. In some experiments, macrophages were exposed for 1 h to titan-like cells (H99 strain, 1:1 ratio) with mAb 18B7. As control, macrophages were also preincubated with the same cells, but without mAb, and also with medium alone (with and without mAb). Then, the wells were washed with fresh medium, and cryptococcal cells of regular size (H99-GFP, grown overnight in Sabouraud at 30 °C) were added at 1:1 ratio for 2 h. After the incubation time, we washed the plates, and separated the macrophages by continuous pipetting. The cell suspensions were washed and suspended in PBS containing 1% FCS. Macrophages were blocked with anti-Fc mAb (2.4G2, BD Biosciences, 5 μg/mL) for 10 min at 4 °C. After that, the macrophages were washed with PBS/FCS, and then incubated with an anti-Mac1 mAb (anti-CD11b/Mac1-PE/Cy7, BioLegend, 1 μg/mL) for 20 min at 4 °C in the dark. Finally, the cells were washed and suspended in 4% p-formaldehyde prepared in PBS. The cells were analyzed in a FACS Canto cytometer (Biosciences, California, EEUU) using FASCDiva software (versión 6.1). The phagocytosis percentage was calculated as followed: (number of PE-Cy7^+^/GFP^+^)/(total number of the PE/PE-Cy7^+^ cells)*100. Experiments were performed in triplicate in three times on different days.

### RNA extraction

RNA extraction was performed using Trizol (TRI Reagent, Sigma Aldrich) with some modifications. After elutriation of the cells, Trizol was added to the samples immediately and maintained in ice. Cells of regular size were broken during 5 minutes with FastPrep^R^ -24 (MP™), alternating 20 seconds beating with 1 minute on ice. Titan cells were disrupted for 10 minutes following the same procedure. The RNAs concentration and quality was determined with a Nanodrop 8000 Spectrophotometer (Thermo scientific). RNA samples (0.1 μg/μL) were treated with DNase using the DNA-free™ kit (Thermo Fisher Scientific). Then, the RNA samples were purified using RNeasy^®^ Mini Kit (Qiagen).

### RNA preparation for RNASeq and transcriptomic data analysis

The total RNA samples (0.5-1 μg) were treated to remove rRNA using Ribo-Zero Magnetic Kit (Epicentre, Illumina, San Diego, CA) according to the manufacturer’s instructions. Then, mRNA was processed for Library preparation using ScriptSeq™ v2 RNA-Seq Library Preparation KIT (Epicentre, Illumina, San Diego, CA). Libraries were quantified using the QuantiFluor® RNA System (Promega) and the quality and average size was determined using an Agilent 2100 Bioanalyzer. An Illumina NextSeq 500 High Output (400 M reads, 1x75 cycles) was using for sequencing. For the results analysis, the FastQ files generated by the equipment were mapped with the software Bowtie2 [90]. Reads were mapped to the genome of strain *C. neoformans* var. *grubbi* H99. The SAM files generated by Bowtie2 were analyzed using the software SeqMonk v0.32.1(Babraham Bioinformatics. The product of the analysis was recovered as *Annotated Probe Report*, which included the coding sequences (overlapping option), and processed in Excel format. The FPKM values for the most differentially expressed genes (>2 fold change, log_2_ >1) were hierarchically clustered using Euclidian distance with Cluster 3.0 program [91]; two clusters of genes with similar expression conditions were identified using k-means clustering. Graphical representation was obtained with the program Java Treeview [92].

To identify functional families of genes that were significantly overrepresented among the overexpressed genes, we selected those that their fold-change difference was at least of two in all the replicas performed. The list of gene was then submitted to FungiFun (https://elbe.hki-jena.de/fungifun/fungifun.php, Hans Knoell Institute).

## Ethics Statement

All the animal procedures were approved by the Bioethical Committee and Animal Welfare of the Instituto de Salud Carlos III (CBA2014_PA51) and of the Comunidad de Madrid (PROEX 330/14) and followed the current Spanish legislation (Real Decreto 53/2013).

## Statistical analysis

Statistical analysis was performed with GraphPad Prism 5. Before comparison among groups, the normality of each simple was assessed using the Kolmogorov-Smirnov test (non-normal distribution when p<0.1). When normal distribution was assumed, differences were estimated using ANOVA and T-Student. For non-parametric distributions, the Kruskal-Wallis and Mann-Whitney tests were used. Statistical significant is highlighted with asterisks in the figures as follows: p>0.5, not significant (ns); p<0.5 and >0.01 (*); p<0.01 and p>0.001 (**); p<0.001 and p>0.0001 (***); p<0.0001 (****).

## Acknowledgements

We want to thank Dr. Guilhem Janbon (Institute Pasteur, Paris, France), Dr. Joseph Heitman (Duke University Medical Center, Durham, North Carolina, United States of America), Dr. Robin May (Birmingham University, UK) and Dr. June Kwon-Chung (NIH, Bethesda, USA), for the gift of strains. I.L. participated in this work in the frame of the master in “Microbiology Applied to Public Health and Research on Infectious Diseases” (Alcalá University and Instituto de Salud Carlos III). We would like to thank Pilar Jiménez Sancho (Genomics Unit, Core Scientific Services, Instituto de Salud Carlos III) for her technical assistance in the RNAseq experiments. We also would like to thank Dr. Manuel Lombardía (Proteomics Facility, National Centre for Biotechnology, Madrid, Spain), for his technical assistance in peptide synthesis.

## Legend for supplemental material

**Supplemental figure 1.** To confirm nuclear analysis, we also used the *C. neoformans* strain SL305 Kn99α NOP1-mCherry [33] that express the nucleolar protein Nop1 tagged with mCherry. In this case, we prepared suspensions of *C. neoformans* cells at 10^4^ and 10^6^ cells/mL in 30 mL of TCM cell. After the incubation period, the cells were collected by centrifugation, fixed with 4% p-formaldehyde and analyzed by flow cytometry using the BD LSRFortessa X-20cytometer (BD, Bioscience). Two populations of titan-like and regular cells were delimited and, in each population, the fluorescence intensity of the NOP1-mCherry protein in 10,000 cells was measured. Data obtained were analyzed with the software BD FACSDiva (BD, Bioscience) and FlowJo 7.6.1 software (Tree Star Inc, Ashland, Oregon). Nuclei of the *C. neoformans* SL305 titan-like and regular cells were also observed by confocal microscopy using a Leica SP5 confocal microscope. In this case, cells were stained with mAb 18B7 conjugated to Alexa-488. A) Titan-like cells; B) cells of regular size. C. Histogram analysis of the NOP1-mCherry fluorescence intensity from cells of regular size (blue) or titan-like cells.

**Supplemental video 1: Cells in Sabouraud medium at 37 °C.** Videos were obtained as described in material and methods. Pictures were taken every 3 minutes, and video was assembled at 12 frames per second, so one second of video corresponds to 36 minutes of real time. Time in each frame reflects the incubation time in TCM.

**Supplemental video 2: Development of titan-like cells.** The yeasts were inoculated in TCM at and incubated overnight at 37 °C with 5% CO_2_. Previous to the start of the video, the cells had been in TCM for 8 h, so the time shown in each frame reflects the total incubation time since the cells were placed in TCM. Videos were assembled as described in supplemental video 1.

**Supplemental video 3: Intracellular features in titan-like cells.** Previous to the start of the video, the cells had been in TCM for 7h 30 min, so the time shown in each frame reflects the total incubation time since the cells were placed in TCM. Video was performed and processed as described above.

**Supplemental video 4. Phagocytosis of titan-like cells by RAW264.7 macrophages**. Videos of the interaction between titan-like cells and macrophages were taken as described in material and methods. Arrow highlights a titan-like cell. Time in the upper right part reflects the time since the beginning of the phagocytosis in minutes.

**Supplemental video 5. Phagocytosis of cells of regular size obtained in TCM by RAW264.7 macrophages**. Cryptococcal cells were incubated in TCM at a cell density of 10^6^ cells/mL, and after one night of incubation at 37 °C with 5% CO_2_, phagocytosis was performed as described in material and methods. Arrow highlights phagocytosis events. Time in the upper right part reflects the time since the beginning of the phagocytosis in minutes.

**Supplemental video 6. Phagocytosis of cells of regular size obtained in Sabouraud by RAW264.7 macrophages**. Cryptococcal cells were incubated in liquid Sabouraud and after one night of incubation at 37 °C with 5% CO_2_, phagocytosis was performed as described in material and methods. Arrow highlights phagocytosis events. Time in the upper right part reflects the time since the beginning of the phagocytosis in minutes.

**Supplemental Table 1.** Elutriation conditions. Conditions used in the process of separation of Titan cells by elutriation. Elutriation system:The centrifugal and drag forces are acting in opposite directions. Unfractionated cells enter the elutriation chamber, size gradient balanced by centrifugal force and counterflow of elutriation fluid keeps cells inside the chamber. Then, increasing flowrate elutes smaller cells first followed by larger ones.

| Elutriation conditions |  |  |  |
| --- | --- | --- | --- |
| Phases | Collection fractions | Centrifugal force (r.p.m) | Counterflow |
| Cell entry | - | 3990 | 50 |
| Cell output |  |  |  |
| **Normal cells** | 1 | 3800 | 50 |
|  | 2 | 3200 | 50 |
|  | 3 | 2700 | 50 |
|  | 4 | 2300 | 50 |
| **Titan cells** | 5 | 1900 | 50 |
|  | 6 | 1500 | 50 |
|  | 7 | 1500 | 80 |
|  | 8 | 1200 | 80 |
|  | 9 | 900 | 80 |
|  | 10 | 500 | 80 |
|  | 11 | Stop | 80 |

**Supplemental Table 2. Transcriptomic analysis of titan-like cells obtained in vitro.**

Gene expression changes in titan-like cells isolated in vitro. Titan-like cells were obtained *in vitro* as described in M&M in two different days, and transcriptomic analysis was performed by RNAseq. The list of genes is present in three different excel worksheets (ALL, UP and DOWN). In ALL, we present the calculated the fold change based on the FPKM values for each gene in each experiment. We selected those than in both replicates were induced by at least two fold (UP worksheet) or repressed by two fold (fold change <0.5, DOWN worksheet).

**Supplemental table 3. Transcriptomic analysis of titan cells isolated from mice.**

Titan and regular cells were isolated from the lung of infected mice as described in M&M and transcriptomic analysis was performed by RNAseq. The experiment was performed in three different days, with three different sets of mice (20 mice each day). The list of genes is present in three different excel worksheets (ALL, UP and DOWN). In ALL, we present the calculated the fold change for each gene in each experiment. In one of the experiments, it was not possible to isolate RNA from one sample of cells of regular size, so the final analysis was performed with three samples from titan cells and two from cells of normal size. To identify genes that behaved similarly in all cases, we calculated in each experiment the fold change of gene expression based on the FPKM values. In the case of the experiment in which we missed the sample of cells of normal size, we calculated the fold change considering as FPKM value the average from the other two experiments. Then, we selected those than in the three replicates were induced by at least two fold (UP worksheet) or repressed by two fold (fold change <0.5, DOWN worksheet). Average fold change was calculated considering the average FPKM from the titan and regular cells samples.

**Supplemental table 4. Genes that were upregulated or repressed in both titan-like and titan cells.** We performed a comparison of the genes that were upregulated or repressed in the two transcriptomic analysis described in supplemental table 2 and supplemental table 3. The list of genes, and the fold change in all the experiments in titan-like (*in vitro*) and titan (*in vivo*) cells are shown in the different worksheets.

